# CRISPR-Cas tools for simultaneous transcription & translation control in bacteria

**DOI:** 10.1101/2023.10.11.561958

**Authors:** Ryan Cardiff, Ian Faulkner, Juliana Beall, James Carothers, Jesse Zalatan

## Abstract

Robust control over gene translation at arbitrary mRNA targets is an outstanding challenge in microbial synthetic biology. The development of tools that can regulate translation will greatly expand our ability to precisely control genes across the genome. In *E. coli*, most genes are contained in multi-gene operons, which are subject to polar effects where targeting one gene for repression leads to silencing of both genes. These effects pose a challenge for independently regulating individual genes in multi-gene operons. Here, we use CRISPR-dCas13 to address this challenge. We find that dCas13-mediated repression exhibits up to 6-fold lower polar effects compared to dCas9. We then show that we can selectively activate single genes in a synthetic multi-gene operon by coupling dCas9 transcriptional activation of an operon with dCas13 translational repression of individual genes within the operon. We also show that dCas13 and dCas9 can be multiplexed for improved biosynthesis of a medically-relevant human milk oligosaccharide. Taken together, our findings suggest that combining transcriptional and translational control can access effects that are difficult to achieve with either mode independently. These combined tools for gene regulation will expand our abilities to precisely engineer bacteria for biotechnology and perform systematic genetic screens.

**Graphical Abstract:** 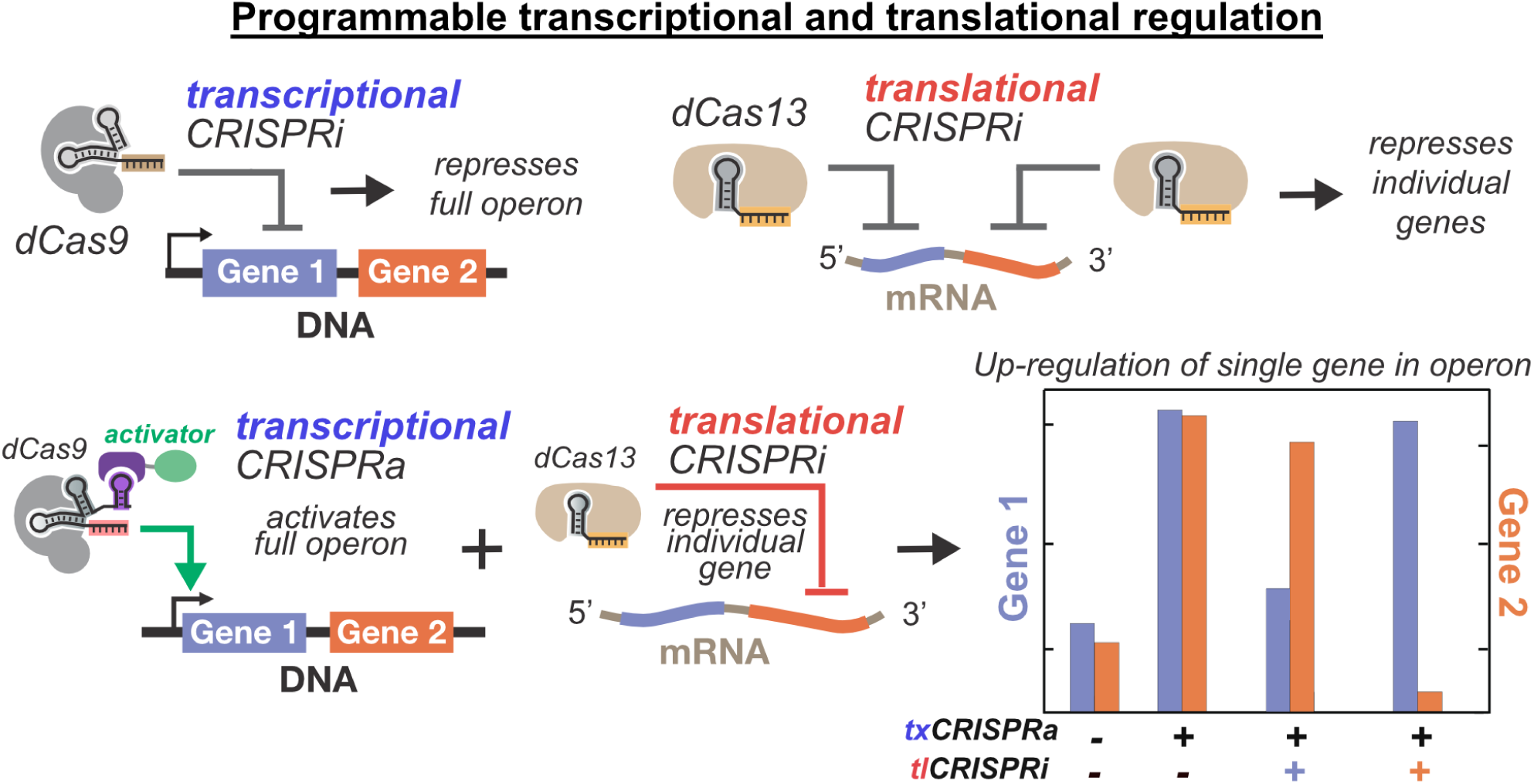

## Introduction

CRISPR-Cas tools have enabled programmable regulation of gene expression in bacteria, allowing control of genomic functions and optimization of biosynthetic production pathways (1–3). To date, most CRISPR-Cas tools that activate, repress, or knockout gene targets in bacteria have been implemented at the DNA level. Despite their enormous potential, these DNA targeting systems have important limitations in their ability to precisely up- or down-regulate individual bacterial gene targets. One major challenge is that targeting one gene in an operon also affects the genes upstream or downstream in the same transcriptional unit (4). In *E. coli*, 68% of all genes are contained in operons containing two or more genes, with some operons containing as many as 14 genes (5, 6). The prevalence of these multi-gene operons poses a challenge for the regulation of individual genes across the genome. Programmable CRISPR-Cas tools that target RNA could potentially overcome this challenge by regulating translation of individual genes.

Programmable RNA targeting with Cas13 has been applied to a variety of applications in eukaryotic and bacterial systems (7–9). In mammalian systems, Cas13 has been used for genome-wide knockdown screens, targeted RNA silencing, and nucleic acid detection (8, 10, 11). The use of catalytically inactive dCas13 fused to an effector domain has also enabled RNA editing and splicing applications (9, 12–14). In *E. coli*, the use of dCas13 for translational repression (CRISPRi) and activation (CRISPRa) of endogenous transcripts has been demonstrated recently (15–18). Despite this progress, research and applications of Cas13 in bacteria are still limited compared to eukaryotic systems.

Here, we characterize dCas13 as a tool for programmable gene regulation in bacteria. Using catalytically inactive dCas13, we first systematically characterize translational silencing (tlCRISPRi) in *E. coli* with a reporter gene assay. We demonstrate tunable repression by varying dCas13 expression levels and guide RNA (gRNA) length, and we characterized sensitivity to target site position and transcript abundance. We then demonstrate that translational-CRISPRi (tlCRISPRi) with dCas13 can preferentially silence specific genes in a multi-gene operon. Although there are still detectable effects on other genes in the operon, these effects are substantially smaller than those seen from transcriptional CRISPRi (txCRISPRi) with dCas9. We then show that we can preferentially activate individual genes in a multi-gene operon by activating the entire operon with transcriptional CRISPRa (txCRISPRa), then downregulating individual genes using tlCRISPRi. Finally, we demonstrate that multiplexed CRISPRa/i tools operating at both the transcriptional and translational levels can improve bioproduction of an important human milk oligosaccharide (HMO) relative to transcriptional control alone.

## Materials & Methods

### Bacterial Strains and Plasmid Constructs

*E. coli* K-12 MG1655 was the parent strain for all flu reporter experiments. JM109, containing a *ΔlacZ* knockout, was used for LNT biosynthesis experiments. CK024 was constructed by integration of Sp.pCas9-dCas9 into MG1655 genome at *arsB* site using the previously described method [Cite Dong-2018]. Strain details/genotypes are described in Table S3. All PCR fragments were amplified with Phusion DNA Polymerase (Thermo-Fisher Scientific) for Infusion Cloning (Takara Bio). Plasmids were transformed into chemically competent NEB Turbo *E. coli* (New England Biolabs) cells, plated on LB-agar, and cultured in LB media supplied with the appropriate antibiotics used in the following concentrations: 100 μg/mL Carbenicillin, 25 μg/mL Chloramphenicol, 30 μg/mL Kanamycin, 100 μg/mL Spectinomycin. All plasmid constructs were confirmed by Sanger sequencing (GENEWIZ). Selected plasmids will be deposited on Addgene (https://www.addgene.org/Jesse_Zalatan/).

All plasmids used in this study are described in Table S1. Relevant DNA sequences are given in the Supplementary Materials. (d)Cas13d and crRNA sequences were cloned from existing plasmids (pXR001 and pXR002) (8). dCas9 plasmids were cloned as described previously (3, 19). Each CRISPR system was expressed in a p15A vector when used alone (Table S1). In Figures 3 and 4, dCas9 and MCP-SoxS (R93A, S101A) (abbreviated MCP-SoxS) were expressed in a pRSF1010 vector to avoid compatibility issues (Table S1) (20). For the multiplexed dCas9/dCas13 experiments in Figure 5, dCas9 was moved to a genome integrated copy (CK024 in Table S3). For the CRISPRa experiment in Figure 5, MCP-SoxS was expressed from the BBa_J23107 promoter on the p15a vector (http://parts.igem.org). For CRISPRa reporter experiments, the guide RNAs for dCas9 and dCas13 were expressed from the strong BBa_J23119 promoter, either in the same plasmid with the Cas-carrying plasmid or in a separate ColE1 plasmid (Table S2). All fluorescent reporters contain the J3 array of CRISPR sites upstream from the minimal promoter. All CRISPRi reporters were expressed from the BBa_J23110 minimal promoter on a psc101** plasmid, unless specified. All CRISPRa reporters and biosynthetic enzymes were expressed from the BBa_J23117 minimal promoter on a pSC101** to provide a greater dynamic range for gene activation.

### Plate Reader Experiments

Single colonies from LB plates were inoculated in 400 μL of EZ-RDM (Teknova) supplemented with the appropriate antibiotics and grown in 96-deep-well plates at 37 °C with shaking overnight 900 rpm on a Heidolph titramax 1000. 100 μL of the overnight culture were transferred into flat, clear-bottomed black 96-well plates (Corning) and the OD_600_ and fluorescence were measured in a Biotek Synergy HTX plate reader. For mRFP1 detection, the excitation wavelength was 540 nm and emission wavelength was 600 nm. For sfGFP detection, the excitation wavelength was 485 nm and emission wavelength was 528 nm. For mBFP detection, the excitation wavelength was 400 nm and emission wavelength was 450 nm. At least three biological replicates were used for all experiments unless stated otherwise.

For kinetic experiments, overnight cultures in stationary phase were subcultured to OD_600_ 0.1 in 200 μL of EZ-RDM (Teknova) supplemented with the appropriate antibiotics. Cultures were grown in flat, clear-bottomed black 96-well plates (Corning) at 37 °C in a Biotek Synergy HTX plate reader set to shake at 1200 RPM. Surrounding wells were filled with 200 μL of water to maintain humidity. The OD_600_ and fluorescence were measured every 30 minutes for 16 hours.

### LNT Production

Single colonies from LB-agar plates were inoculated in 2 mL EZ-RDM (Teknova) with 10 g/L glucose, 2 g/L lactose and supplemented with appropriate antibiotics. Cultures were grown in 14 mL glass culture tubes at 37 °C and shaking for 48 h. 500 μL of supernatant from each culture were loaded onto 10 kDa microcentrifuge filters (Millipore) and spun for 20 min at 14000 rcf.

Samples were analyzed via Agilent 6530 LC/Q-TOF in negative mode using a Poroshell 120 C18 150 mm column. A standard curve was prepared by spiking known amounts of LNT (Elicityl) into supernatants derived from cultures of MG1655 or JM109 *E. coli* transformed with empty vectors. For LC/MS, the aqueous phase was water with 0.1% formic acid and the organic phase was acetonitrile with 0.1% formic acid. The % aqueous/organic gradient was run as follows: hold at 3/97 for 1 minute, 10/90 over 1 minute, 40/60 over 3 minutes, hold 40/60 for 2 minutes, then return to 3/97 over 2.5 minutes. The flow rate was held at 0.3 mL/min.

## Results

### Optimizing tlCRISPRi repression in *E. coli*

To repress translation in *E. coli*, we need to deliver dCas13 and a CRISPR RNA (crRNA) that targets the mRNA of a gene of interest. We used the catalytically inactive version of RfxCas13d (referred to as dCas13 from here), which was shown to have greater knockdown efficiency than Cas13a and Cas13b in mammalian cells (8). We provided crRNAs targeting either an mRFP fluorescent reporter gene or an off-target control. When expressed without dCas13, the crRNA targeting mRFP gave 1.4-fold repression compared to the parental strain (Figure 1A), an effect which has previously been reported for Cas13 systems (15). This crRNA-dependent effect may be due to direct binding of the mRNA target, forming a dsRNA structure and interfering with mRNA function (21, 22). When dCas13 was expressed with the crRNA, the overall tlCRISPRi effect improved to 2.9-fold (Figure 1A). We also tested dCas13d against another dCas13 isoform, dCas13a from *Leptotrichia buccalis* (23), to confirm that dCas13d remained the most effective in *E. coli*. We found that dCas13d was more active, consistent with previous results in mammalian cells (8) (Figure S1). Together, these results show that dCas13 can be used with a crRNA for programmable translational repression.

**Figure 1.**
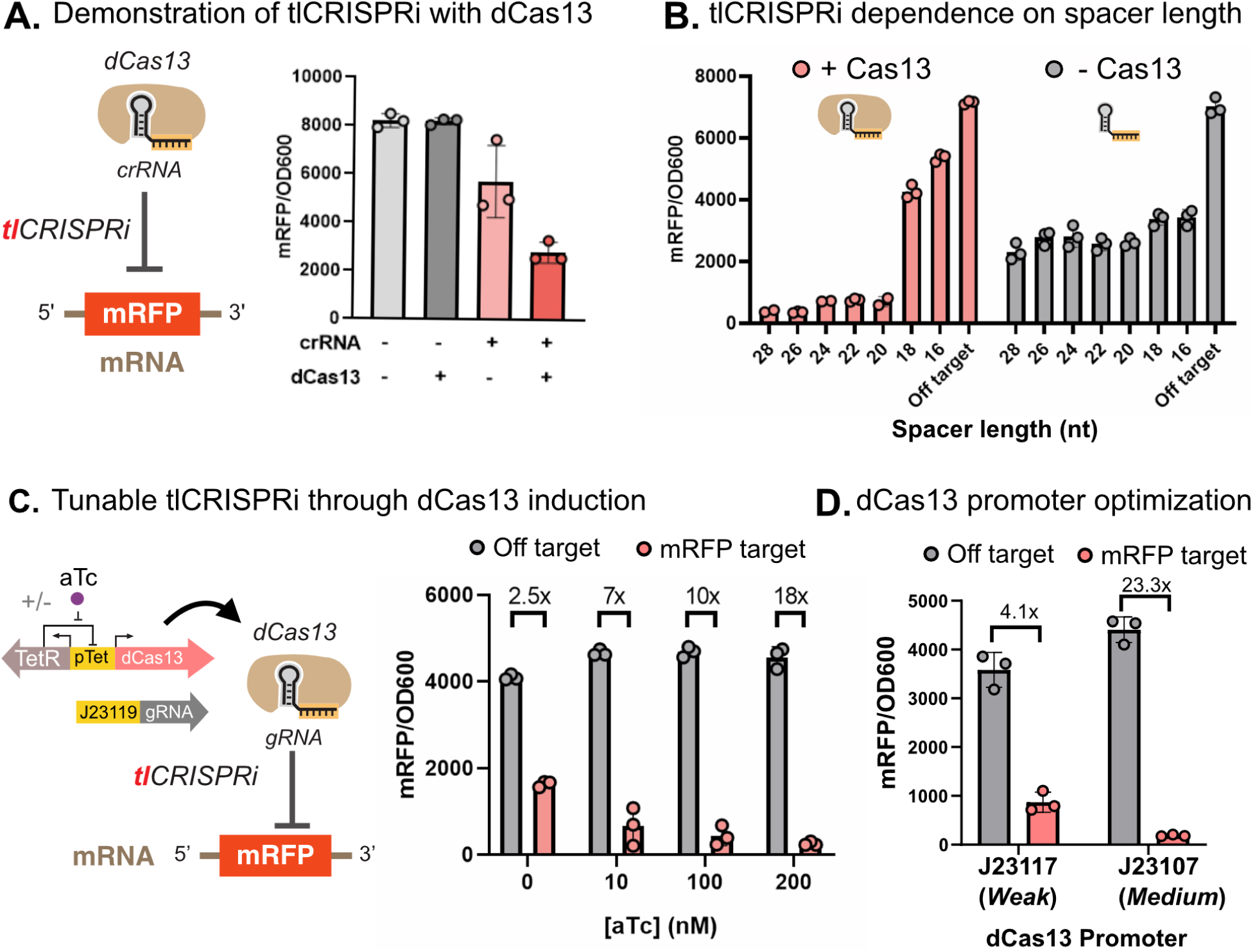
tlCRISPRi represses gene expression in *E. coli*. A) Effect on reporter gene expression with dCas13 alone, crRNA alone, or both dCas13 and crRNA.The crRNA *rfp2* targets the ORF of the mRFP mRNA transcript (Table S2). B) Reporter gene expression versus crRNA spacer length. The spacer is the region complementary to the target mRNA. Reporter gene expression levels were measured both in the presence and absence of dCas13. The *rfp2* crRNA was tested with spacer lengths from 16-28 by truncating 2 nt at a time from the 5’ end. C) Tuning tlCRISPRi activity by titrating dCas13 expression levels. dCas13 was expressed with an inducible pTet promoter and the crRNA was expressed from the constitutive J23119 promoter. Reporter gene expression levels were measured with 0, 10, 100 and 200 nM aTc concentrations. D) tlCRISPRi with dCas13 expression from weak (J23117) or medium (J23107) strength constitutive promoters. For all panels, values represent the mean ± standard deviation for at least three biological replicates.

To further investigate the effects of the crRNA alone on translation, we measured tlCRISPRi for a fluorescent reporter as a function of crRNA spacer sequence length both in the presence and absence of dCas13. crRNAs were truncated 2 bp at a time from an initial spacer length of 28 nt. Spacers were truncated from the 5’ end, although previous work has shown that truncations from either the 5’ or 3’ end follow similar qualitative trends (11). We found that the crRNA-alone effect results in roughly a 2-fold repression when crRNA length varied between 16 - 28 nt (Figure 1B), similar to the effects observed in the previous experiment (Figure 1A). With dCas13 present, we observed repression effects ranging from 1.5-fold up to 18-fold, with the strongest repression from longer crRNAs (≥20 nt) (Figure 1B). Together, these results demonstrate that the both dCas13 and the crRNA are required for maximal repression, the natural crRNA length of 26-28 nt is optimal for dCas13-mediated repression, and varying the spacer length from 16-28 nt does not change the crRNA-alone effects.

Having demonstrated simple tlCRISPRi, we sought to further optimize the dCas13 expression levels. Because dCas9 has been reported to be toxic to *E. coli* cells at high expression levels (24, 25), we cloned dCas13 under the inducible Tet promoter. aTc titrations showed that *E. coli* is tolerant to relatively high dCas13 concentrations (Figure S2). We found that tlCRISPRi repression improves up to 200 nM aTc induction (Figure 1C). We then cloned dCas13 under weak and medium strength constitutive promoters and observed a similar trend where stronger expression gave greater repression (Figure 1D). We also observed that catalytically-inactive dCas13d was 7-fold more effective at RFP repression than wild-type, nuclease-active Cas13d (Figure S3). This result is surprising because target mRNA cleavage should further reduce translation beyond any tlCRISPRi binding effects. To our knowledge, this direct comparison of CRISPRi efficiency between Cas13 and dCas13 has not been made previously. More effective repression from dCas13 may be due to collateral RNA cleavage activity of the nuclease-active form leading to additional toxicity to the cell (26, 27), although we cannot exclude the possibility that there are differences in protein expression levels for Cas13 and dCas13.

To better understand crRNA targeting rules for tlCRISPRi, we designed four crRNAs targeting different positions of the mRFP transcript. We found that as the distance from the 5’ end of the mRNA transcript increased, the tlCRISPRi efficiency decreased (Figure 2A). This observation is in agreement with previous findings that have found targeting the unstructured ribosome binding site (RBS) to be most effective for dCas13-mediated knockdowns (15). This result is also consistent with similar distance-dependent effects in other CRISPR systems (28).

**Figure 2.**
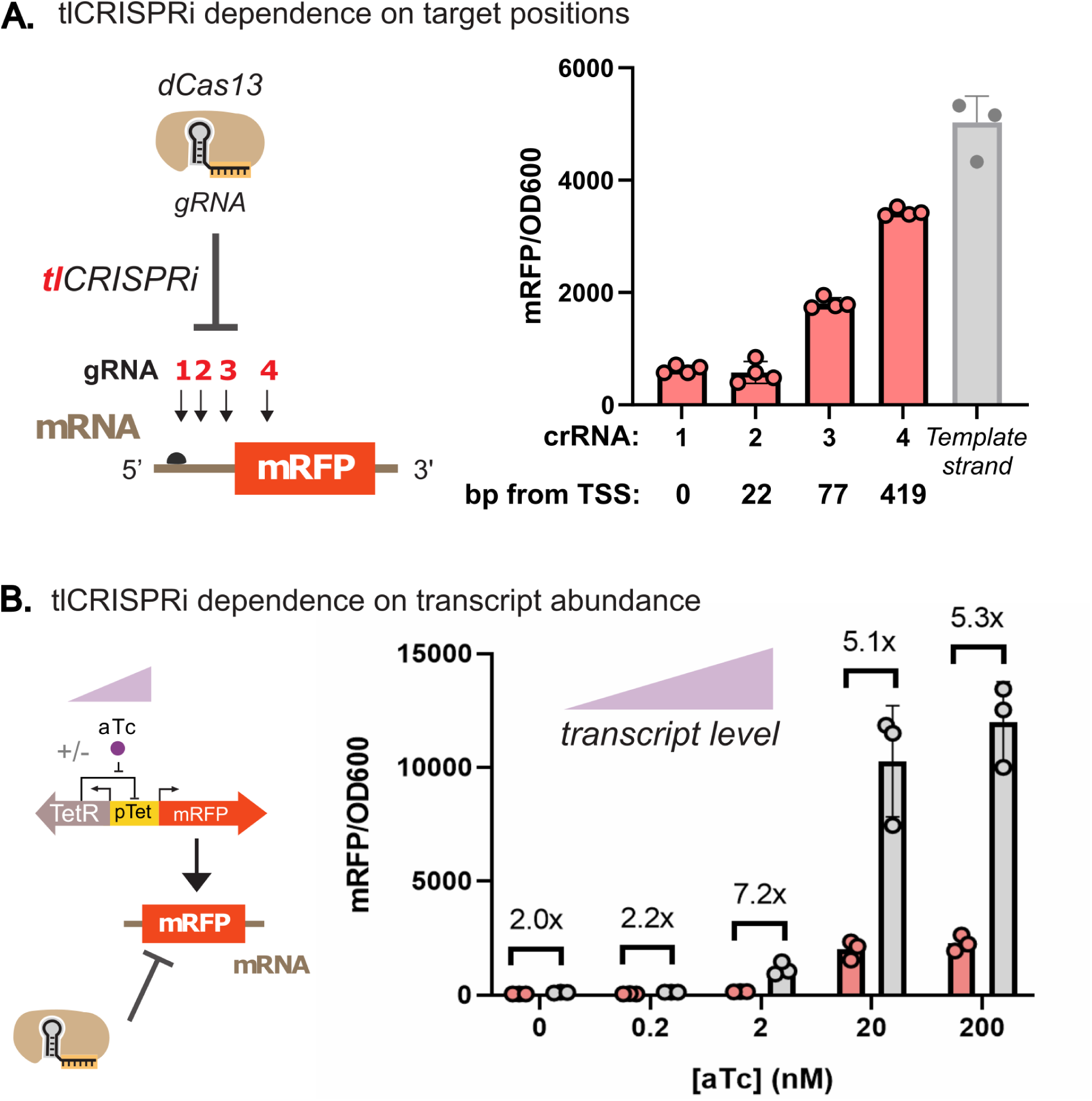
tlCRISPRi dependence on mRNA target site position and transcript abundance. A) Reporter gene expression versus tlCRISPRi target position on the mRNA. *rfp1* targets the unstructured RBS. *rfp2* targets the 3’ end of the RBS and beginning of the ORF. *rfp3* and *rfp4* target the middle of the ORF. The off-target control targets the template strand, which is not transcribed into mRNA. B) tlCRISPRi effects on reporter gene expression versus target mRNA transcript expression level. mRFP was expressed from the pTet promoter and induced with 0, 0.2, 2, 20, and 200 mM aTc. dCas13 and crRNA were constitutively expressed. For all panels, values represent the mean ± standard deviation for at least three biological replicates.

In addition to target position, we anticipated that tlCRISPRi efficiency would depend on the mRNA abundance in the cell. We used an pTet-inducible mRFP reporter to measure tlCRISPRi while titrating aTc, effectively titrating the number of target mRNA copies in the cell. tlCRISPRi is most effective at 2 nM aTc. At higher aTc concentrations and correspondingly higher target mRNA abundance, repression may be limited by the amount of CRISPR complexes in the cell (Figure 2B) (29). At lower aTc concentrations and correspondingly lower transcript abundance, the expression level approaches the lower limit of detection, which limits the detectable dynamic range of repression.

### tlCRISPRi has smaller polar effects than txCRISPRi

While transcriptional control systems act on entire operons, RNA targeting with dCas13 may enable independent regulation of single genes in multi-gene operons. The use of dCas13 may greatly improve our ability to regulate bacterial functions, as a large fraction of genes are contained in multi-gene operons. To test whether dCas13 can independently repress one gene while leaving the other unperturbed, we built a synthetic reporter constitutively expressing sfGFP and mRFP in a single transcriptional unit. The dual operon reporter uses a strong constitutive promoter (J23110) to express the sfGFP operon reading frame (ORF) followed by the mRFP ORF, each translated from a separate synthetic Bujard RBS (30).

We first tested the dual operon reporter with the DNA-targeting dCas9, which we expected would produce strong polar effects where targeting one gene for repression leads to silencing of both genes (31). As expected, dCas9 exhibited strong polar effects when targeting either gene in the operon (Figure 3A). We quantified polar effects by comparing the ratio of repression between the target gene and the other gene in the operon. Targeting the upstream sfGFP gene gave 15-fold repression of sfGFP and 34-fold repression of mRFP, representing a 2.3-fold weaker repression of sfGFP than the downstream mRFP. Targeting the downstream mRFP gene gave 32-fold repression of mRFP and 5.7-fold repression of sfGFP, an overall 5.6-fold greater repression of mRFP than sfGFP (Figure 3A). The weaker polar effect on sfGFP when downstream mRFP is targeted may be due to incomplete mRNA transcripts that still allow for some translation of the first gene in the operon.

**Figure 3.**
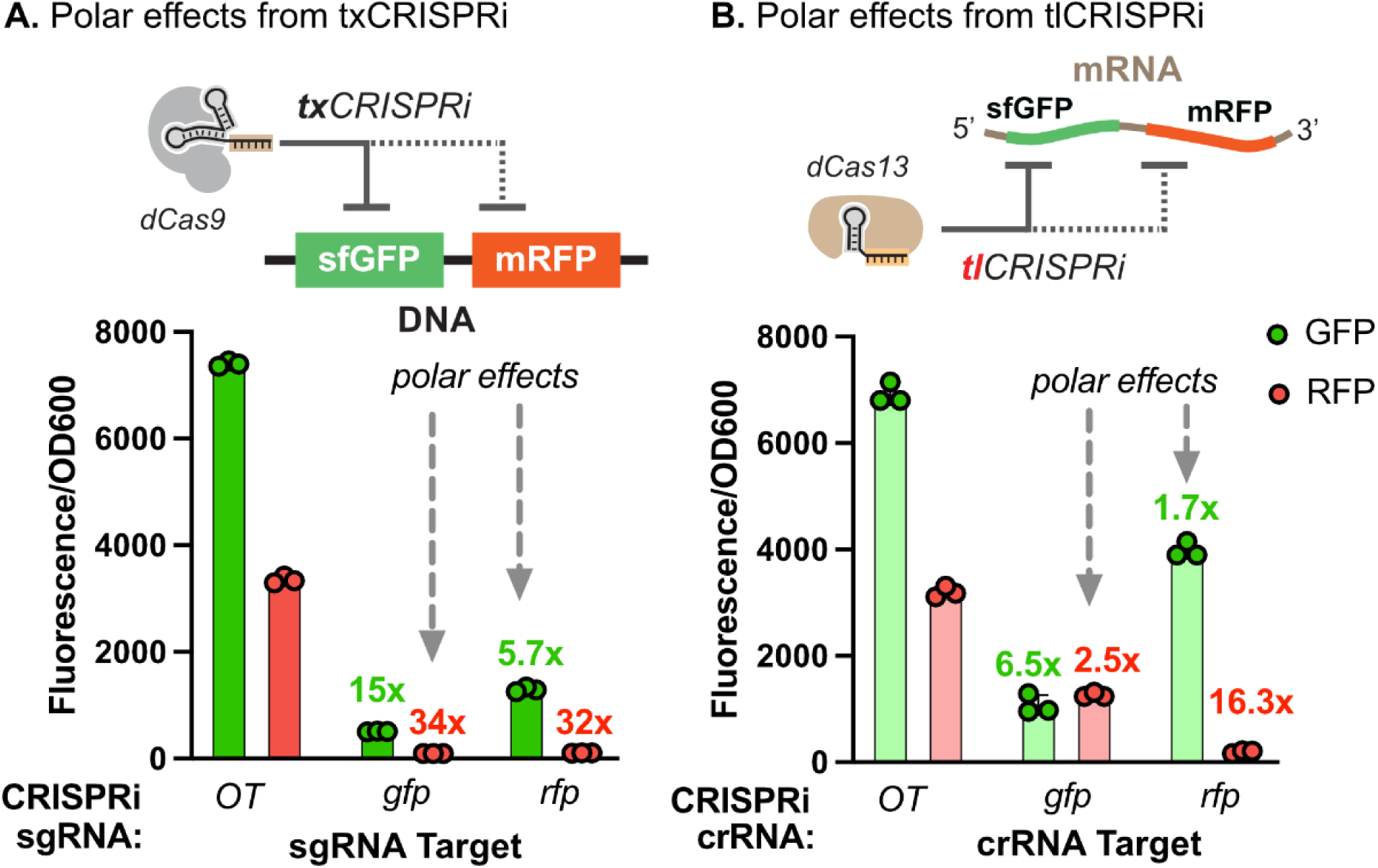
CRISPRi polar effects for multi-gene operons. A) Polar effects from txCRISPRi with dCas9-mediated targeting of DNA to repress transcription. A synthetic multi-gene sfGFP-mRFP reporter transcript was expressed from the J3_J23110 promoter (see Methods). Each gene has a separate copy of the Bujard RBS. Polar effects on reporter gene expression were measured by comparing the ratio of repression between the target gene and the other gene in the operon. Fluorescent reporter levels were measured with dCas9 sgRNAs for sfGFP (*NT1)*, mRFP (*rr2)*, or an off-target (OT) (*AAV)* sequence (Table S2). B) Polar effects from tlCRISPRi with dCas13-mediated targeting of mRNA to repress translation. The reporter gene was the same as described above in (A). Fluorescent reporter levels were measured with dCas13 crRNAs for sfGFP (*gfp2*), mRFP (*rfp2)*, or an off-target (OT) (*AAV)* sequence (Table S2). For all panels, values represent the mean ± standard deviation for at least three biological replicates.

We then used dCas13 to test tlCRISPRi on each gene in the dual reporter operon. dCas13 was expressed from the medium strength J23107 promoter to produce a large tlCRISPRi effect based on our previous results (Figure 1D). Compared to transcriptional repression with dCas9, we observed reduced polar effects (Figure 3B). Targeting mRFP, the second ORF in the operon, gave 16.3-fold repression of mRFP, but only 1.7-fold repression sfGFP (Figure 3B). There was a 10-fold greater repression of mRFP than sfGFP, showing that we can target a downstream gene in an operon with modest effects on upstream genes. Targeting sfGFP gave 6.5-fold repression of sfGFP and 2.5-fold repression of mRFP, representing a 2.6-fold greater repression of sfGFP than mRFP. This result indicates that there is a larger polar effect when targeting the upstream gene compared to the downstream gene. For both genes, the observed polar effects with dCas13 were less than that of dCas9 repression (Figure S4).

### Multiplexed dCas9/dCas13 tools improve independent regulation of multi-gene operons

We next tested whether we could activate a subset of genes in a multi-gene operon by activating transcription of the entire operon and selectively repressing translation for undesired genes. As a proof of concept, we envisioned using a previously described dCas9 txCRISPRa system (19, 32) to activate a two-gene operon together with dCas13 as a translational repressor. We hypothesized dCas13 to repress one gene in the operon while leaving the other activated.

To investigate simultaneous txCRISPRa and tlCRISPRi, we modified the dual sfGFP/mRFP operon used in previous experiments. We replaced the strong constitutive promoter with a weak promoter that could be activated with dCas9. Using the txCRISPRa system alone with dCas9 targeting the promoter and no crRNA for dCas13 targeting, sfGFP and mRFP were activated 10 and 16-fold, respectively (Figure 4A). When a crRNA was introduced to target mRFP with dCas13, we observe that mRFP returns to basal levels while sfGFP remains activated. This result shows that we can specifically activate the first gene in a multi-gene operon by upregulating the entire operon, then using tlCRISPRi to repress downstream genes. However, when we tried to repress sfGFP with dCas13, we observed low levels of repression that were equivalent for sfGFP and mRFP (Figure 4A). This result was unexpected because we previously observed preferential sfGFP repression when targeting sfGFP in the constitutively-expressed dual reporter, although by a relatively modest factor of 2.6-fold (Figure 3B). Taken together, our results suggest that independently repressing a downstream gene is effective, but independently repressing an upstream gene remains challenging.

**Figure 4.**
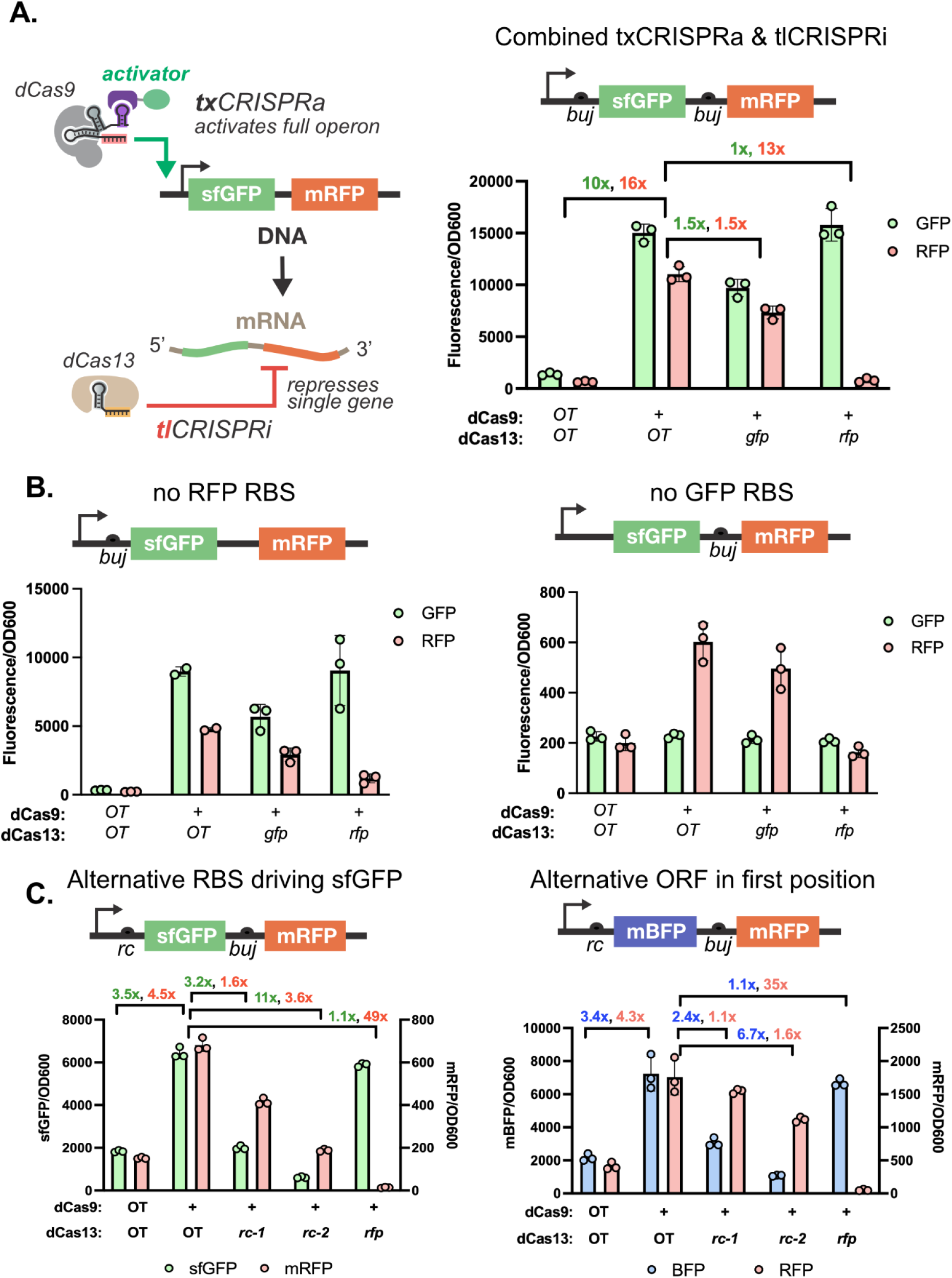
Multiplexed txCRISPRa/tlCRISPRi in multi-gene operons. A) Combined txCRISPRa and tlCRISPRi for independent gene activation in a multi-gene operon. For txCRISPRa, dCas9 targets the −81 position of the J3_J23117 promoter and recruits a transcriptional activator to activate the operon (19). For tlCRISPRi, dCas13 targets an individual gene with a crRNA targeting either sfGFP or mRFP. The dCas13 off-target (OT) sample indicates the fold activation from txCRISPRa without any tlCRISPRi repression. Fold repression from tlCRISPRi compares the dCas13 off-target and on-target crRNAs, always with the on-target J306 scRNA for txCRISPRa. B) Combined txCRISPRa/tlCRISPRi as described in (A) using a modified reporter lacking an RBS for either mRFP (left) or sfGFP (right) to investigate translational coupling. C) Combined txCRISPRa/tlCRISPRi as described in (A) using the *rc* RBS at the first gene in the operon instead of the Bujard RBS. The first gene in the operon is either sfGFP (left) or mBFP (right). Two new crRNA sequences were designed to target the *rc* RBS (Table S2). For all panels, values represent the mean ± standard deviation for at least three biological replicates.

The strong polar effect on the downstream mRFP gene when targeting the upstream sfGFP could be due to translational coupling of the two genes (33, 34). There are many examples of translational coupling between genes in natural systems. Translational coupling can occur if ribosomes translating the upstream gene are recruited to the downstream gene, either through stop codon read-through or increased local concentration near the downstream RBS (33, 35). Alternatively, ribosome-mediated mRNA unfolding can disrupt secondary structures that inhibit translation initiation at downstream genes (36–38). Thus, if translation of the downstream gene depends wholly or in part on translation of the upstream gene, polar effects on downstream genes with tlCRISPRi may be difficult to avoid.

To test whether translational coupling could explain the observed polar effects, we constructed modified reporters where the RBS had been removed for either sfGFP or mRFP and repeated the multiplexed txCRISPRa/tlCRISPRi experiment. The reporter lacking a RBS for the upstream sfGFP showed greatly reduced mRFP levels (Figure 4B) compared to the reporter with the sfGFP RBS (Figure 4A). This result suggests that a significant amount of the observed mRFP expression is from translational coupling to the upstream sfGFP. However, the mRFP decrease could also result from decreased mRNA transcript stability that can occur when the RBS is removed from the 5’ end (39). We found stronger evidence for translational coupling by removing the RBS from the downstream mRFP. With this reporter, we observed that mRFP was still upregulated when the operon was activated with dCas9 (Figure 4B), again suggesting that there is significant translational coupling from the upstream sfGFP to the downstream mRFP.

To test if independent repression of the upstream gene in an operon is possible, we need a reporter with minimal translational coupling between genes. Ribosome read-through is thought to be dependent on the rate of translational initiation at the upstream gene (33, 34). We therefore tested an alternative *rc* RBS for the upstream sfGFP that has lower sfGFP expression when activated by txCRISPRa (Figure S5). We designed two crRNA targets for this RBS to repress sfGFP translation, *rc-1* and *rc-2*. With these two crRNAs, we observed 3.2- and 11-fold repression of sfGFP, while the effects on the downstream mRFP were 1.6- and 3.6-fold, respectively. These effects correspond to 2.0- or 3.1-fold preferential repression of sfGFP than mRFP (Figure 4C, left), in contrast to equivalent repression of sfGFP and mRFP seen with the original *buj* RBS at sfGFP (Figure 4A). Thus, it is possible to selectively repress the upstream gene, although the preferential effect is relatively modest. Notably, the preferential dCas13 effects with this txCRISPRa-activated reporter are similar to the 2.6-fold preferential repression we observed with the original constitutive reporter (Figure 3B). In the constitutive reporter, the initial sfGFP and mRFP levels were similar to the levels produced by txCRISPRa-mediated activation with the *rc* RBS (Figure 4C, left). Thus, the extent of translational coupling, and consequently the possibility of independently repressing the first gene in an operon, may be dependent on the absolute expression level.

We then assessed whether the sfGFP mRNA sequence context could be contributing to the observed translational coupling and polar effects. We replaced the sfGFP reporter with mBFP (mTagBFP2, blue fluorescent protein) under the same *rc* RBS. Qualitatively, we observed similar effects with either mBFP or sfGFP in the upstream position. Targeting the downstream mRFP with dCas13 strongly silences mRFP by ~35-fold with only a 1.1-fold effect on mBFP (Figure 4C, right). Targeting the RBS of the upstream mBFP with the *rc-1* and *rc-2* crRNAs produced 2.4- and 6.7-fold repression of mBFP, while the effects on the downstream mRFP were 1.1- and 1.6-fold, respectively. These effects correspond to 2.2- and 4.2-fold higher repression of mBFP than the downstream mRFP, respectively (Figure 4C, right). These effects are qualitatively similar to those observed with sfGFP in the upstream position (2.0- and 3.1-fold, respectively) but the quantitative differences suggest that transcript sequence context can contribute to polar effects.

One potential disadvantage of the combined txCRISPRa/tlCRISPRi approach is that there may be a growth burden associated with expressing two CRISPR systems. To test whether the cell could support the two systems operating simultaneously, we measured the growth kinetics of a cell expressing either the txCRISPRa, tlCRISPRi, or both txCRISPRa/tlCRISPRi machineries. We found that expressing dCas13 and dCas9 together in the cell imposed no detectable additional burden to the cell compared to expressing the CRISPRa machinery alone (Figure S6).

### Multiplexed dCas9/dCas13 metabolic engineering improves LNT bioproduction

To test whether a multiplexed dCas9 and dCas13 system could be effective for metabolic engineering, we used bacterial lacto-N-tetraose (LNT) production as a model system. LNT is a human milk oligosaccharide (HMO) that is important for infant immune development and microbiome health (40, 41). To produce LNT in *E. coli*, two heterologous enzymes, LgtA and *Cv*GalT, can be combined with overexpression of the LacY lactose permease (42, 43). LacY imports lactose into the cell, where LgtA, a β-1,3-*N*-acetylglucosaminyltransferase from *N. meningitidis,* produces the intermediate lacto-*N*-triose II (LNT II). CvGalT is a β-1,3-galactosyltransferase from *C. violaceum* that converts LNT II to the final LNT product. Knocking out endogenous β-galactosidase activity (encoded by the *lacZ* gene) is also necessary to prevent cleavage of the lactose feedstock into its constituent monosaccharides glucose and galactose (43).

Because LNT production requires up- and down-regulation of multiple enzymes, CRISPR-Cas-mediated gene regulation can be useful for tuning and optimizing gene expression levels. We previously used a combinatorial txCRISPRa library to optimize pathway enzyme levels in a *lacZ* knockout strain [*in preparation*] (Figure 5A). Here, we attempted to further optimize the system by using CRISPRi to repress endogenous genes that divert flux away from LNT. Specifically, we targeted the *ugd, wecB,* and *nagB* genes. The enzymes from these genes consume the metabolites UDP-*N*-acetylglucosamine and UDP-galactose, which are used by LgtA and CvGalT during LNT production. Previous work has shown that knocking these genes out can improve LNT production by factors ranging from 20 - 350% (44–46). However, when we activated the LNT pathway enzymes with txCRISPRa and repressed *ugd, wecB,* or *nagB* using txCRISPRi, we observed that the knockdowns produced no increase in LNT production relative to the strain with txCRISPRa alone (Figure 5B).

**Figure 5.**
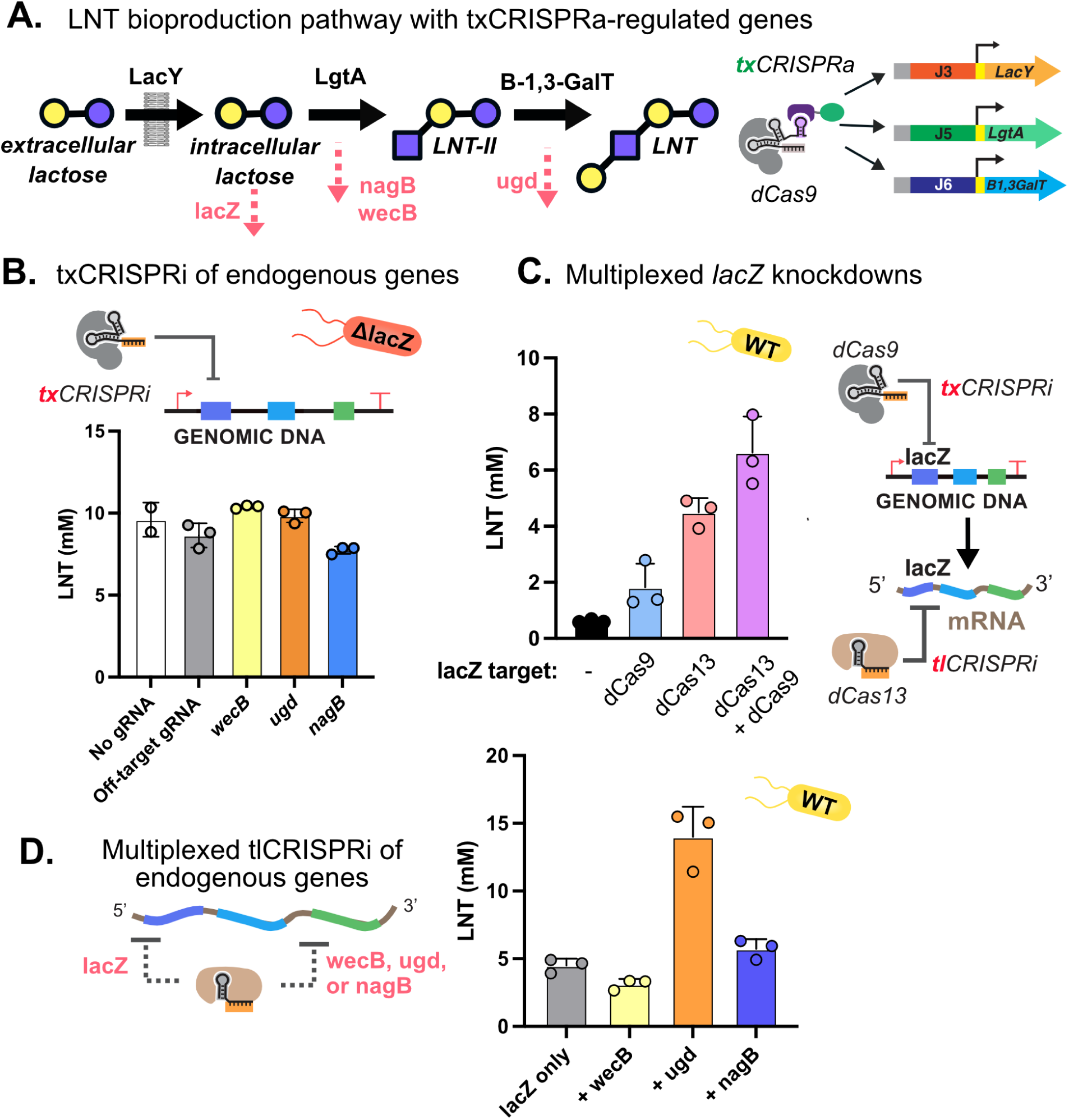
Multiplexed txCRISPRa/tlCRISPRi for improved metabolic engineering. A) Pathway schematic for LNT bioproduction in *E. coli* with CRISPRa-regulated genes. LNT is produced from lactose using three plasmid-expressed enzymes. The heterologous pathway is weakly expressed from synthetic promoters and activated using txCRISPRa. The pathway consists of *lacY* from *E. coli*, *lgtA* from *Neisseria meningitidis,* and *B3GalT* from *Chromobacterium violaceum.* Endogenous genes that divert flux away from the LNT pathway, including *wecB, ugd,* and *nagB*, are shown in red. These genes are targets for repression to increase LNT flux. B) LNT titers with different endogenous genes targeted for repression with txCRISPRi. Experiments were conducted in the JM109 *E. coli* strain, which contains a *lacZ* deletion to prevent lactose metabolism. C) LNT titers with txCRISPRi and tlCRISPRi individual and multiplexed knockdowns of *lacZ.* Experiments were conducted in the CK024 *E. coli* strain, which contains wild type (WT), unmodified *lacZ* and genome integrated dCas9 (Table S3). D) LNT titers for multiplexed dCas13 knockdowns of *lacZ* and other endogenous gene targets. Experiments were conducted in the CK024 *E. coli* strain (Table S3). For all panels, values represent the mean ± standard deviation for three biological replicates.

We next tested whether dCas13-mediated tlCRISPRi knockdowns could improve LNT production. We targeted *lacZ* for knockdown in an *E. coli* strain containing the native copy to demonstrate we could produce a sufficient knockdown for LNT production. We compared *lacZ* knockdowns from dCas13 and dCas9 to the strain containing the *lacZ* knockout. We found that the dCas13 knockdown gave 4.5 ± 0.5 mM LNT compared to 1.8 ± 0.8 mM from the dCas9 knockdown (Figure 5C). However, both knockdowns produced less LNT than the 8.6 ± 0.7 mM produced from the *lacZ* knockout (Figure 5B). This result could be due to the CRISPR perturbations not producing a strong enough knockdown, or from increased cellular burden from the operation of the CRISPR systems. To test this, we multiplexed dCas13 and dCas9 knockdowns of *lacZ* and observed 6.6 ± 1.3 mM of LNT production (Figure 5C). This result suggests that the knockdowns from each system were not as strong as the knockout, and that greater knockdowns can be achieved by combining perturbations.

One potential advantage of CRISPR regulation is the opportunity for rapid combinatorial multiplexing. In the strain containing the dCas13 *lacZ* crRNA, we introduced a second crRNA targeting either *ugd, nagB,* or *wecB*. While dCas9 knockdowns of these targets all gave lower or similar levels of LNT, the dCas13 knockdown of *ugd* gave a 3-fold improvement in LNT production, from 4.5 ± 0.5 mM to 14 ± 2.2 mM (Figure 5D). The dCas13 *nagB* knockdown also gave a modest increase in LNT production to 5.7 ± 0.7 mM (Figure 5D). The combined *lacZ* and *ugd* dCas13-mediated knockdown strain was the highest producing strain, yielding a 46% improvement over the *lacZ* knockout strain with no translational regulation.

## Discussion

CRISPR-Cas technologies have enabled a range of applications in bacterial systems, including gene editing, biosensing, and bioproduction (3, 47–50). The ability to combine orthogonal CRISPR-Cas systems with different regulatory properties can expand the possibilities for precise gene regulation and information processing. Additionally, combining orthogonal CRISPR-Cas systems may overcome limitations of gRNA competition to allow regulation of more genes in the genome simultaneously. The ability to increase the upper limit of regulatable gene targets in a single metabolic engineering program will be an important consideration for future efforts (51).

Here, we show that multiplexed DNA- and RNA-targeting dCas9 and dCas13 CRISPR-Cas systems can improve bioproduction in bacteria. These results emphasize that different methods of regulation can yield different and difficult to predict effects when regulating endogenous genes. The use of dCas13, either alone or in addition to dCas9, may offer several advantages for metabolic engineering. Previous HMO metabolic engineering efforts have shown *lacZ* knockout to be critical for effective production (44–46). Here, we showed that multiplexed knockdowns of *lacZ* with dCas9 and dCas13 improved LNT production relative to either dCas9 or dCas13 alone. Multiplexed knockdowns of additional genes can yield further improvements to LNT production to 14 mM. To our knowledge, this titer is the highest achieved to date in shake flask conditions. We have also shown that dCas9 and dCas13 can be combined to overcome polar effects in multi-gene transcriptional operons. The ability to independently regulate individual genes in a multi-gene operon depends on sequence context, expression level, and target position within the operon, but we have shown that polar effects can be partially mitigated through screening different crRNA targets (Figure 4C). Genome-wide tlCRISPRi screens could help to further elucidate sequence and context dependent rules that determine the polar effects at multi-gene operons.

Many targets for knockdown may not produce the desired effects when repressed with dCas9 due to tight regulation at the transcriptional level. For example, *nagB* is known to be highly regulated due to its relevance for sugar carbon utilization (52, 53). When targeted with dCas9, the *nagB* knockdown resulted in a significantly lower LNT production than the other knockdown strains (Figure 5B). We then showed when nagB was targeted with dCas13 that LNT production increased from 4.5 to 5.7 mM (Figure 5D). dCas13 may also provide an advantage over dCas9 when targeting genes in the same operon as an essential gene. Systematic comparisons of dCas9 and dCas13 knockdowns may uncover other contexts in which translational control is preferred over transcriptional regulation.

Greater investigation into Cas13 systems has the potential to improve our ability to precisely regulate individual genes at the translational level. Dynamic control of dCas13 activity could enable improved biosynthesis, molecular recording, biosensing, and biotherapeutic applications (54–57). Such a system could be constructed by regulating dCas13 expression with an inducible or conditionally-responsive promoter. However, because there is weak basal repression from the crRNA alone (Figure 1B), tight control of crRNA expression may be necessary for maximal dynamic range.

The ability to activate gene expression using dCas13 could also be useful for a wide range of bacterial engineering applications. Our results here suggest that dCas13 mediated activation (tlCRISPRa) could offer advantages over DNA-targeting dCas9 (txCRISPRa) by enabling control of individual genes in multi-gene operons. txCRISPRa is also subject to relatively stringent target requirements that have limited its generalizability to arbitrary gene targets (19, 58, 59). If tlCRISPRa with dCas13 is subject to less stringent or even simply different target requirements, it will expand the range of accessible gene targets in bacteria. Previous work in *E. coli* has shown that dCas13 can be fused to an effector domain for translational activation (tlCRISPRa) (15). Still, robust tlCRISPRa on arbitrary mRNA targets remains to be demonstrated (15, 18).

Moving forward, the ability to use dCas13 to independently regulate individual genes across bacterial genomes along with multiplexed-regulation using dCas9 may greatly expand the possibilities for basic biology research and applied biotechnology. dCas13 has already proven a useful tool for studying gene essentiality in mammalian and bacterial systems (11, 27). Genome-wide dCas13 screens may reveal novel insights into phenotypes from individual ORF perturbations that would be missed from dCas9 knockdown screens that act at the operon-level. Multi-gene operons are ubiquitous across nature. However, tools to study individual genes within these operons are limited. Improved control over multi-gene operons will expand our understanding of individual gene functions. In addition to essentiality studies, genome-wide tlCRISPRi screens can inform strain design for a variety of applications. The realization of dCas13 tools in medically and industrially relevant microbes will assist the development of living therapeutics, bioproduction, and *in situ* bioremediation.

## Data sharing plans

All data for this work are available in the Supplementary Information file.

## Acknowledgements

We thank members of the Carothers and Zalatan groups, especially Cholpisit (Ice) Kiattisewee, for advice, materials, and comments on the manuscript.

