## Supplementary Information for "CRISPR-Cas tools for simultaneous transcription & translation control in bacteria"

1: Molecular Engineering & Sciences Institute
and Center for Synthetic Biology
University of Washington
Seattle, WA 98195
United States

2: Department of Chemical Engineering
University of Washington
Seattle, WA 98195
United States

3: Department of Chemistry
University of Washington
Seattle, WA 98195
United States

*Corresponding authors


206-221-4902


206-543-1670

**Supplementary Figures**

**Figure S1:** Effect of tlCRISPRi on mRFP expression using dCas13d or dCas13a
**Figure S2:** Endpoint $OD_{600}$ for dCas13 and Cas13
**Figure S3:** Reporter gene expression for tlCRISPRi using nuclease-inactive dCas13 and nuclease-active Cas13
**Figure S4:** Fold repression ratio for two fluorescent reporters in an operon targeted by both dCas9 and dCas13
**Figure S5:** Gene expression for txCRISPRa of two reporter constructs
**Figure S6:** Growth kinetics measured from cells containing dual-CRISPR systems

**Supplementary Tables**

**Table S1:** Plasmids used in this study
**Table S2:** gRNA sequences
**Table S3:** Strains used in this study
**Table S4:** DNA Sequences

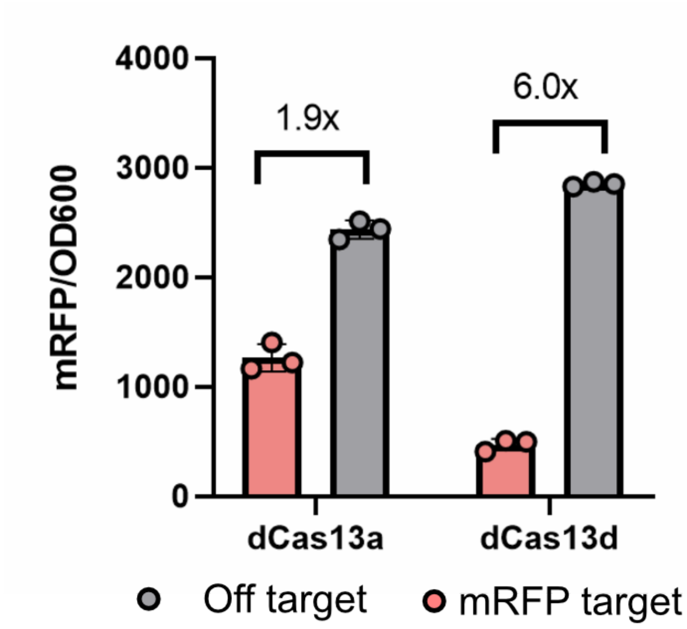


Figure S1. Effect of tlCRISPRi on mRFP expression using dCas13d or dCas13a. The crRNAs *rfp2* and *AAV* were used as on- and off-targets, respectively (Table S2). dCas13 variants were induced with 200 nM aTc. Values represent the mean ± standard deviation for at least three biological replicates.

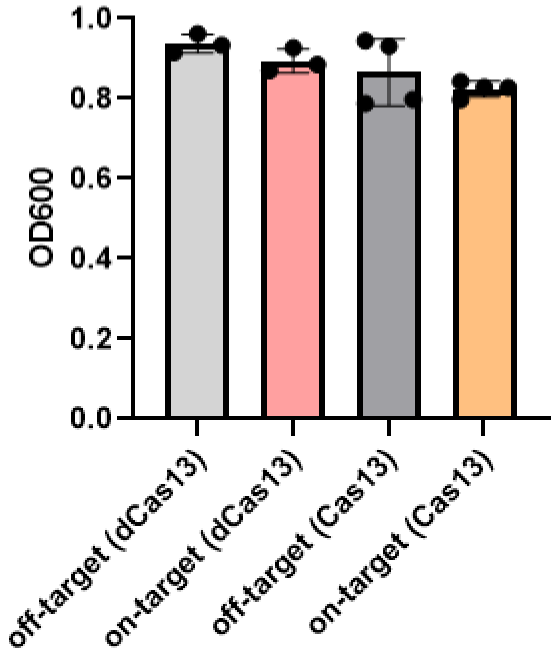


Figure S2. Endpoint $OD_{600}$ values for *E. coli* cultures were measured after 18 hours with dCas13 and Cas13 induced with 200 nM aTc. The crRNAs *rfp2* and *AAV* were used as on- and off-targets, respectively, targeting a mRFP reporter gene (Table S2). Values represent the mean ± standard deviation for at least three biological replicates.

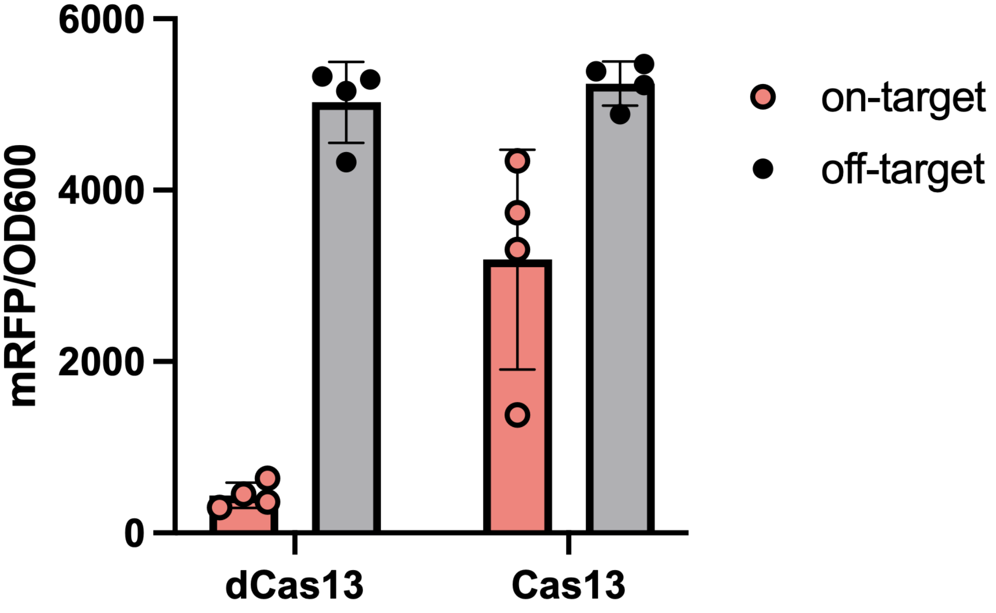


Figure S3. Reporter gene expression for tlCRISPRi using nuclease-inactive dCas13 and nuclease-active Cas13. The mRFP reporter was constitutively expressed from the J3_J23110 promoter. The crRNAs *rfp2* and *AAV* are used as on- and off-targets, respectively (Table S2). dCas13 and Cas13 were induced with 200 nM aTc. Values represent the mean ± standard deviation for at least three biological replicates.

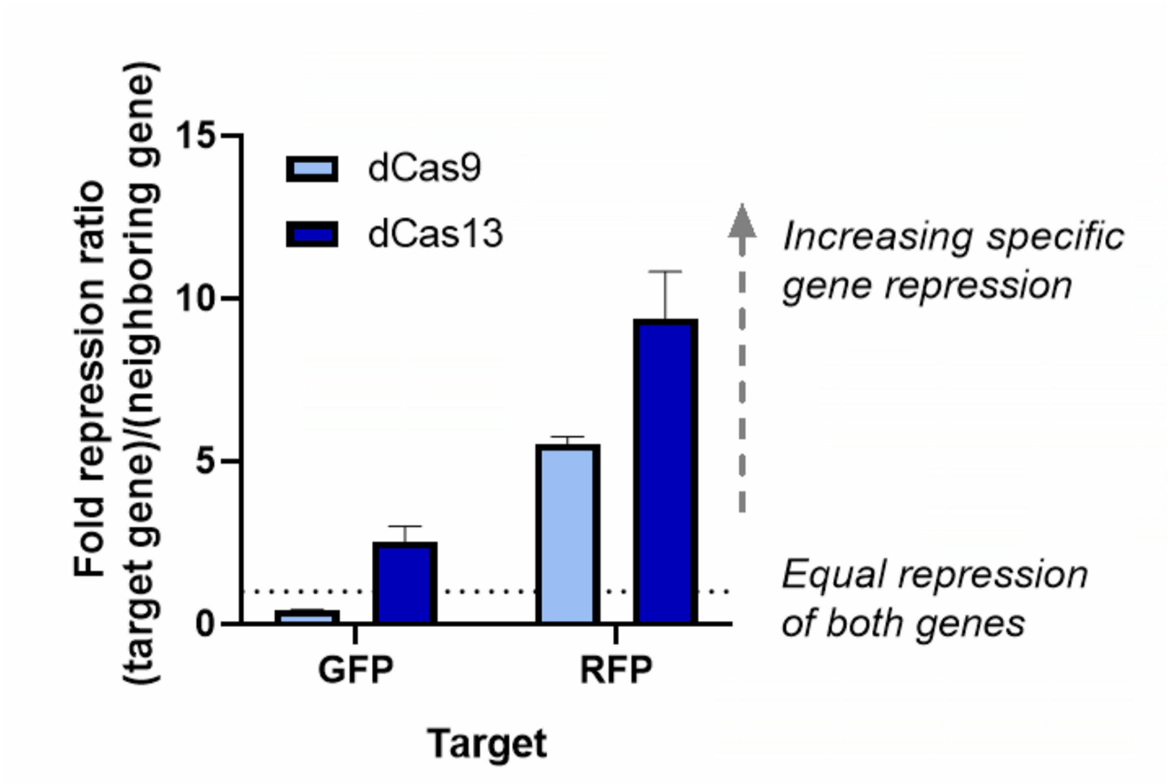


Figure S4. Fold repression ratio for two fluorescent reporters in an operon targeted by both dCas9 and dCas13. For each reporter, the ratio of fold repression for the target gene to fold repression of the neighboring operon gene is shown on the Y-axis as a measure of polar effects. CRISPRi polar effects are compared between dCas9 (light blue) and dCas13 (dark blue). The reporter operon consists of sfGFP followed by mRFP, expressed from a strong constitutive promoter (J3_J23110) and separate copies of the Bujard RBS. Accompanying raw data and fold repression are shown in Figure 3. For dCas9, the sgRNA targets *rr2* and *NT1* were used to target mRFP and sfGFP respectively, and fold repression values were calculated relative to cells expressing an off target sgRNA for *AAV* (Table S2). For dCas13, the crRNA targets *rfp2* and *gfp2* were used to target mRFP and sfGFP, and fold repression values were calculated relative to cells expressing an off target crRNA for *AAV* (Table S2).

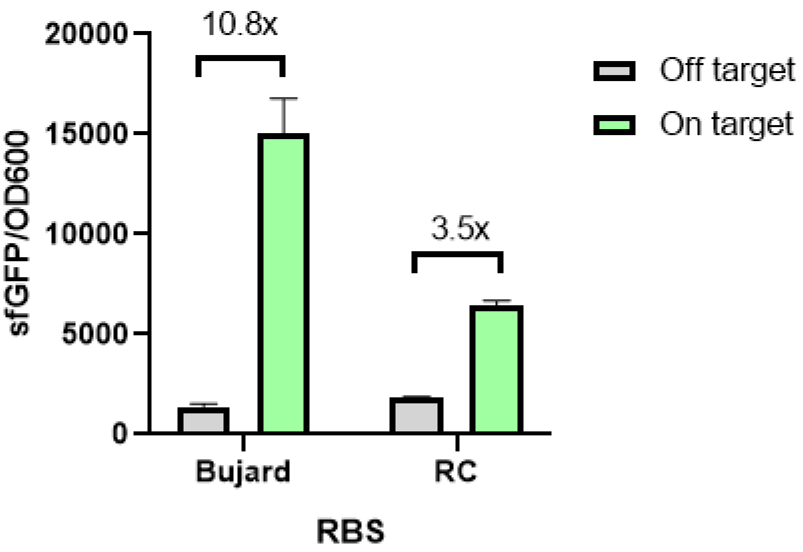


Figure S5. Gene expression for txCRISPRa of two reporter constructs. txCRISPRa was performed on multi-gene operons containing either the Bujard (left) and *rc* (right) synthetic RBSs driving expression of sfGFP, the first gene in the operon. The reporter uses the weak constitutive promoter J3_J23117, which allows for improved activation range over the previously used J23110 minimal promoter. The on-target gRNA used for dCas9 was J306 targeting -81 bp from the TSS, and the off-target gRNA is AAV (Table S2). Values represent the mean ± standard deviation for at least three biological replicates.

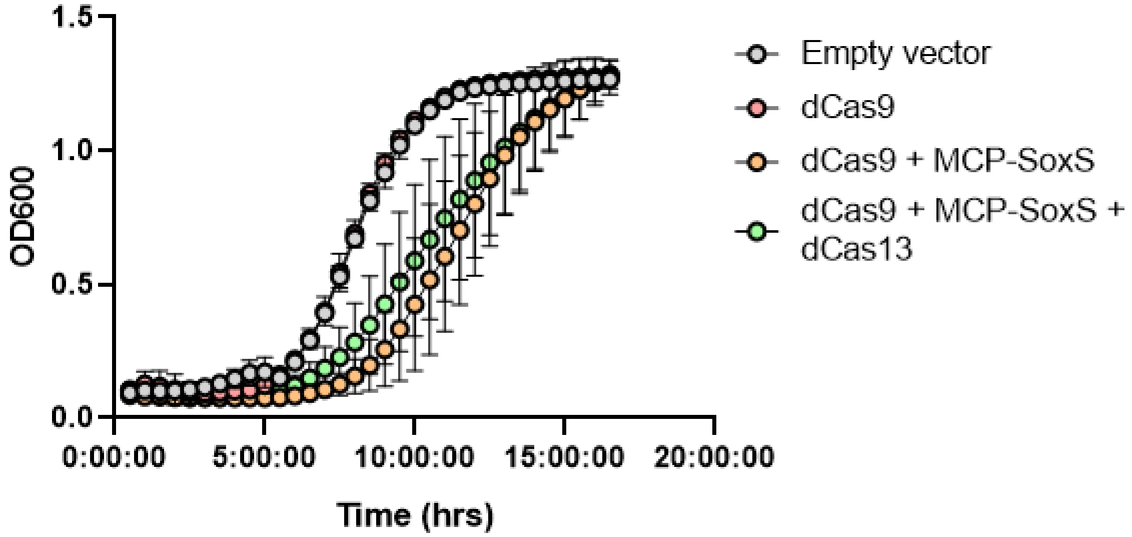


Figure S6. Growth kinetics measured from cells containing dual-CRISPR systems. For the cells containing empty vector and dCas9 + MCP-SoxS, the JM109 strain was used (Table S3) containing the plasmids pJF043 or pJF234, respectively (Table S1). For the cells containing dCas9 alone or dCas9, MCP-SoxS, and dCas13, the CK24 strain was used (Table S3). For the dCas9, MCP-SoxS, and dCas13 condition, the plasmid pRC172 was used (Table S1). Growth conditions are described in the Materials & Methods. Values represent the mean ± standard deviation for at least three biological replicates.

Table S1. Plasmids used in this study

| Plasmid | Marker | Replicon | Promoter | RBS | Gene | Terminator | Reference |
|---|---|---|---|---|---|---|---|
| pJF143.J3_J23110 | Amp | psc101** | J3_J23110 | buj | mRFP | dblTerm | (31), Figures 1 and 2 |
| pRC055 | Chlor | p15a | pTet | rc-2 | dRxCas13d | dblTerm | This work, Figure 1 |
| pRC056 | Chlor | p15a | pTet | rc-2 | RxCas13d | dblTerm | This work, Figure S1 |
| pRC057.crRNA | Chlor | p15a | pTet<br>J23119 | rc-2<br>- | dRxCas13d<br>crRNA | dblTerm<br>ECK120033736 | This work, Figure 1 |
| pRC058.crRNA | Chlor | p15a | pTet<br>J23119 | rc-2<br>- | RxCas13d<br>crRNA | dblTerm<br>ECK120033736 | This work, Figure S1 |
| pRC055.117 | Chlor | p15a | J23117 | rc-2 | dRxCas13d | dblTerm | This work, Figure 1 |
| pRC055.107 | Chlor | p15a | J23107 | rc-2 | dRxCas13d | dblTerm | This work, Figure 1 |
| pRC065.crRNA | Chlor | p15a | J23107<br>J23119 | rc-2<br>- | dRxCas13d<br>crRNA | dblTerm<br>ECK120033736 | This work, Figure 1 |
| pRC065d.crRNA | Chlor | p15a | J23107<br>J23119 | rc-2<br>- | dRxCas13d<br>crRNA | dblTerm<br>ECK120033736 | This work, Figure 1 |
| pRC073.crRNA | Chlor | p15a | J23119 | - | crRNA | ECK120033736 | This work, Figure 1 |
| pRC146 | Kan | ColE1 | J23119 | - | crRNA | ECK120033736 | This work, Figure 1 |
| pRC072 | Amp | psc101** | J3_23110 | buj, buj | sfGFP, mRFP | dblTerm | This work, Figure 3 |
| pJB003 | Amp | psc101** | J3_J23117 | buj, buj | sfGFP, mRFP | dblTerm | This work, Figure 4 |

| | | | | | | | |
|---|---|---|---|---|---|---|---|
| pRC239 | Amp | psc101** | J3_J23117 | rc, buj | sfGFP,<br>mRFP | dblTerm | This work,<br>Figure 4 |
| pRC241 | Amp | psc101** | J3_J23117 | rc, buj | mBFP,<br>mRFP | dblTerm | This work,<br>Figure 4 |
| pJF083 | Amp | psc101** | pTet | buj | mRFP | dblTerm | This work,<br>Figure 2 |
| pCK580 | Chlor | psc101** | pTet | rc-2 | dCas13a | dblTerm | This work,<br>Figure S2 |
| pCK513.gRNA | Spec | pRSF1010 | J23107,<br>J23107,<br>J23119 | Buj,<br>Buj,<br>- | dCas9,<br>MCP-SoxS,<br>scRNA | dblTerm,<br>BBa_B1002,<br>ECK120033736 | This work,<br>Figure 3 |
| pJF234 | Chlor | p15a | Sp-Pdcas9,<br>J23107<br>J23105,<br>J23105,<br>J23105 | Native RBS,<br>Buj,<br>-<br>-<br>- | dCas9,<br>MCP-SoxS,<br>J306<br>J506<br>J606 | dblTerm,<br>BBa_B1002,<br>ECK120033736 (x3) | This work,<br>Figure 5 |
| pJF234.gRNA | Chlor | p15a | Sp-Pdcas9,<br>J23107<br>J23105,<br>J23105,<br>J23105,<br>J23110 | Native RBS,<br>Buj,<br>-<br>-<br>-<br>- | dCas9,<br>MCP-SoxS,<br>J306,<br>J506,<br>J606,<br>dCas9 gRNA | dblTerm,<br>BBa_B1002,<br>ECK120033736 (x3) | This work,<br>Figure 5 |
| pIDF96C | Amp | psc101** | J3,<br>J5,<br>J6 | RBS-A,<br>RBS-B,<br>RBS-C | lacY,<br>lgtA,<br>galT | ECK120033737,<br>ECK120010818,<br>ECK120015440 | This work,<br>Figure 5 |
| pRC172 | Chlor | p15a | J23107,<br>J23107<br>J23105,<br>J23105,<br>J23105 | rc2,<br>buj,<br>-<br>-<br>- | dCas13,<br>MCP-SoxS,<br>J306<br>J506<br>J606 | dblTerm,<br>BBa_B1002,<br>ECK120033736 (x3) | This work,<br>Figure 5 |
| pRC182 | Chlor | p15a | J23107, | rc2, | dCas13, | dblTerm, | This work, |

| | | | | | | | |
|---|---|---|---|---|---|---|---|
| | | | J23107<br>J23105,<br>J23105,<br>J23105,<br>J23110 | buj,<br>-<br>-<br>-<br>- | MCP-SoxS,<br>J306,<br>J506,<br>J606,<br>dCas13<br>gRNA | BBa_B100<br>2,<br>ECK12003<br>3736 (x4) | Figure 5 |
| pRC188 | Chlor | p15a | J23107<br>J23105,<br>J23105,<br>J23105 | buj,<br>-<br>-<br>- | MCP-SoxS,<br>J306<br>J506<br>J606 | BBa_B100<br>2,<br>ECK12003<br>3736 (x3) | This work,<br>Figure 5 |
| pRC189 | Chlor | p15a | J23107<br>J23105,<br>J23105,<br>J23105,<br>J23110 | buj,<br>-<br>-<br>-<br>- | MCP-SoxS,<br>J306,<br>J506,<br>J606,<br>dCas13<br>gRNA | dblTerm,<br>BBa_B100<br>2,<br>ECK12003<br>3736 (x4) | This work,<br>Figure 5 |
| pJF043 | Chlor | p15a | - | - | - | - | This work,<br>Figure S6 |

Table S2. gRNA sequences

| **gRNA** | **DNA Sequence** | **CRISPR System** | **Target** |
|---|---|---|---|
| *rfp1* | TGGTACCTTTCTCC<br>TCTTTAATGAATTC | dCas13 | RFP |
| *rfp2* | AgAACGTCTTCGCT<br>ACTCGCCATGGTAC | dCas13 | RFP |
| *rfp3* | ctcgtgaccgttaa<br>cggaaccttccata | dCas13 | RFP |
| *rfp4* | tttttttctgcata<br>accggaccgtcgga | dCas13 | RFP |
| *rfp5* | tgcttacaaaaccg<br>acatcaaactggac | dCas13 | RFP |
| *gfp2* | ATGAGCAAAGGAGA<br>AGAACTTTTCACTG | dCas13 | GFP |
| *rc1* | GTGCTCCCTCGTGA | dCas13 | *rc* RBS (expressing |

| | | | |
|---|---|---|---|
| | GAACTTTTTCGAAC | | either sfGFP or mBFP) |
| *rc2* | ATAAAACCTCCTTACTCTGTGCTCCCTC | dCas13 | *rc* RBS (expressing either sfGFP or mBFP) |
| *lacZ-9* | TTCTCCGTGGGAACAAACGG | dCas9 | lacZ |
| *nagB-9* | GTTCACGCAACCTTCAACGG | dCas9 | nagB |
| *ugd-9* | TACATAGCCAGTACCGGAAA | dCas9 | ugd |
| *wecB-9* | ACAAACGGTAAATACTCCTG | dCas9 | wecB |
| *lacZ-13* | cggccagtgaatccgtaatcatggtcat | dCas13 | lacZ |
| *nagB-13* | ggatcagtctcattattcacctcaataa | dCas13 | nagB |
| *ugd-13* | agccagtaccggaaatggtgattttcat | dCas13 | ugd |
| *wecB-13* | aatacagtcagtactttcacatcgattc | dCas13 | wecB |
| *J306* | TTGTGTCCAGAACGCTCCGT | dCas9 | -81bp from the TSS of synthetic J3 promoter |
| *J506* | AGCAGCATGAGCAGCATTGA | dCas9 | -81bp from the TSS of synthetic J5 promoter |
| *J606* | GTCGCAGTCGCGCGAGCACT | dCas9 | -81bp from the TSS of synthetic J6 promoter |
| *AAV* | GGGGCCACTAGGGACAGGAT | dCas13/dCas9 | Off-target |
| *rr2* | TGGAACCGTACTGGAACTGC | dCas9 | RFP |

| | | | |
|---|---|---|---|
| *NT1* | CATCTAATTCAACA AGAATT | dCas9 | GFP |

Table S3. Strains used in this study

| Strain | Description | Genotype |
|---|---|---|
| MG1655 | Wildtype *E. coli* strain | F- λ- ilvG- rfb-50 rph-1 |
| CK24 | Integrated dCas9 used for combined dCas9 + dCas13 experiments | MG1655_arsB::Sp.pCas9-dCas9 |
| JM109 | *lacZ* knockout used for dCas9-only LNT experiments | *end*A1, *rec*A1, *gyr*A96, *thi*, *hsd*R17 ($r_k^-$, $m_k^+$), *rel*A1, *sup*E44, Δ( *lac-pro*AB), [F´ *tra*D36, *pro*AB, *lac*I$^q$ZΔM15] |

Table S4. DNA Sequences

**Reporters**

>J3_J23110_**buj**_sfGFP_**buj**_mRFP_dblTerm

AGCATTTGCGATCATTCACGCAGCGCTTATTCAGTTGCTCACTGCGATGTCATAATCATCGCTA
CGAGCTGTGAAAGATGCATAAAGCTCGTACGACGCGTTCGCTCGTCTCCTCACTTCTCCTACGG
AGCGTTCTGGACACAACGTCGTCTTGAAGTTGCGATTATAGAtttacggctagctcagtcctag
gtacaatgctagc**GAATTCATTAAAGAGGAGAAAGGTACC**ATGAGCAAAGGAGAAGAACTTTTC
ACTGGAGTTGTCCCAATTCTTGTTGAATTAGATGGTGATGTTAATGGGCACAAATTTTCTGTCC
GTGGAGAGGGTGAAGGTGATGCTACAAACGGAAAACTCACCCTTAAATTTATTTGCACTACTGG
AAAACTACCTGTTCCGTGGCCAACACTTGTCACTACTCTGACCTATGGTGTTCAATGCTTTTCC
CGTTATCCGGATCACATGAAACGGCATGACTTTTTCAAGAGTGCCATGCCCGAAGGTTATGTAC
AGGAACGCACTATATCTTTCAAAGATGACGGGACCTACAAGACGCGTGCTGAAGTCAAGTTTGA
AGGTGATACCCTTGTTAATCGTATCGAGTTAAAGGGTATTGATTTTAAAGAAGATGGAAACATT
CTTGGACACAAACTCGAGTACAACTTTAACTCACACAATGTATACATCACGGCAGACAAACAAA
AGAATGGAATCAAAGCTAACTTCAAAATTCGCCACAACGTTGAAGATGGTTCCGTTCAACTAGC
AGACCATTATCAACAAAATACTCCAATTGGCGATGGCCCTGTCCTTTTACCAGACAACCATTAC
CTGTCGACACAATCTGTCCTTTCGAAAGATCCCAACGAAAAGCGTGACCACATGGTCCTTCTTG

AGTTTGTAACTGCTGCTGGGATTACACATGGCATGGATGAGCTCTACAAAtaaggatcc**GAATT**
**CATTAAAGAGGAGAAAGGTACC**ATGGCGAGTAGCGAAGACGTTATCAAAGAGTTCATgcgtttc
aaagttcgtatggaaggttccgttaacggtcacgagttcgaaatcgaaggtgaaggtgaaggtc
gtccgtacgaaggtacccagaccgctaaactgaaagttaccaaaggtggtccgctgccgttcgc
ttgggacatcctgtccccgcagttccagtacggttccaaagcttacgttaaacacccggctgac
atcccggactacctgaaactgtccttcccggaaggtttcaaatgggaacgtgttatgaacttcg
aagacggtggtgttgttaccgttacccaggactcctccctgcaagacggtgagttcatctacaa
agttaaactgcgtggtaccaacttcccgtccgacggtccggttatgcagaaaaaaaccatgggt
tgggaagcttccaccgaacgtatgtacccggaagacggtgctctgaaaggtgaaatcaaaatgc
gtctgaaactgaaagacggtggtcactacgacgctgaagttaaaaccacctacatggctaaaaa
accggttcagctgccgggtgcttacaaaaccgacatcaaactggacatcacctcccacaacgaa
gactacaccatcgttgaacagtacgaacgtgctgaaggtcgtcactccaccggtgcttaaggat
ccaaactcgagtaaggatctCCAGGCATCAAATAAAACGAAAGGCTCAGTCGAAAGACTGGGCC
TTTCGTTTTATCTGTTGTTTGTCGGTGAACGCTCTCTACTAGAGTCACACTGGCTCACCTTCGG
GTGGGCCTTTCTGCGTTTATA

**dCas9**

>Sp-PdCas9_dCas9_dblTerm
TTACGAAATCATCCTGTGGAGCTTAGTAGGTTTAGCAAGATGGCAGCGCCTAAATGTAGAATGA
TAAAAGGATTAAGAGATTAATTTCCCTAAAAATGATAAAACAAGCGTTTTGAAAGCGCTTGTTT
TTTTGGTTTGCAGTCAGAGTAGAATAGAAGTATCAAAAAAAGCACCGACTCGGTGCCACTTTTT
CAAGTTGATAACGGACTAGCCTTATTTTAACTTGCTATGCTGTTTTGAATGGTTCCAACAAGAT
TATTTTATAACTTTTATAACAAATAATCAAGGAGAAATTCAAAGAAATTTATCAGCCATAAAAC
AATACTTAATACTATAGAATGATAACAAAATAAACTACTTTTTAAAAGAATTTTGTGTTATAAT
CTATTTATTATTAAGTATTGGGTAATATTTTTTGAAGAGATATTTTGAAAAAGAAAAATTAAAG
CATATTAAACTAATTTCGGAGGTCATTAAAACTATTATTGAAATCATCAAACTCATTATGGATT
TAATTTAAACTTTTTATTTTAGGAGGCAAAAATGGATAAGAAATACTCAATAGGCTTAGcTATC
GGCACAAATAGCGTCGGATGGGCGGTGATCACTGATGAATATAAGGTTCCGTCTAAAAAGTTCA
AGGTTCTGGGAAATACAGACCGCCACAGTATCAAAAAAAATCTTATAGGGGCTCTTTTATTTGA
CAGTGGAGAGACAGCGGAAGCGACTCGTCTCAAACGGACAGCTCGTAGAAGGTATACACGTCGG
AAGAATCGTATTTGTTATCTACAGGAGATTTTTTCAAATGAGATGGCGAAAGTAGATGATAGTT
TCTTTCATCGACTTGAAGAGTCTTTTTTGGTGGAAGAAGACAAGAAGCATGAACGTCATCCTAT
TTTTGGAAATATAGTAGATGAAGTTGCTTATCATGAGAAATATCCAACTATCTATCATCTGCGA
AAAAAATTGGTAGATTCTACTGATAAAGCGGATTTGCGCTTAATCTATTTGGCCTTAGCGCATA
TGATTAAGTTTCGTGGTCATTTTTTGATTGAGGGAGATTTAAATCCTGATAATAGTGATGTGGA
CAAACTATTTATCCAGTTGGTACAAACCTACAATCAATTATTTGAAGAAAACCCTATTAACGCA
AGTGGAGTAGATGCTAAAGCGATTCTTTCTGCACGATTGAGTAAATCAAGACGATTAGAAAATC
TCATTGCTCAGCTCCCCGGTGAGAAGAAAAATGGCTTATTTGGGAATCTCATTGCTTTGTCATT
GGGTTTGACCCCTAATTTTAAATCAAATTTTGATTTGGCAGAAGATGCTAAATTACAGCTTTCA
AAAGATACTTACGATGATGATTTAGATAATTTATTGGCGCAAATTGGAGATCAATATGCTGATT
TGTTTTTGGCAGCTAAGAATTTATCAGATGCTATTTTACTTTCAGATATCCTAAGAGTAAATAC

TGAAATAACTAAGGCTCCCCTATCAGCTTCAATGATTAAACGCTACGATGAACATCATCAAGAC
TTGACTCTTTTAAAAGCTTTAGTTCGACAACAACTTCCAGAAAAGTATAAAGAAATCTTTTTTG
ATCAATCAAAAAACGGATATGCAGGTTATATTGATGGGGGAGCTAGCCAAGAAGAATTTTATAA
ATTTATCAAACCAATTTTAGAAAAAATGGATGGTACTGAGGAATTATTGGTGAAACTAAATCGT
GAAGATTTGCTGCGCAAGCAACGGACCTTTGACAACGGCTCTATTCCCCATCAAATTCACTTGG
GTGAGCTGCATGCTATTTTGAGAAGACAAGAAGACTTTTATCCATTTTTAAAAGACAATCGTGA
GAAGATTGAAAAAATCTTGACTTTTCGAATTCCTTATTATGTTGGTCCATTGGCGCGTGGCAAT
AGTCGTTTTGCATGGATGACTCGGAAGTCTGAAGAAACAATTACCCCATGGAATTTTGAAGAAG
TTGTCGATAAAGGTGCTTCAGCTCAATCATTTATTGAACGCATGACAAACTTTGATAAAAATCT
TCCAAATGAAAAAGTACTACCAAAACATAGTTTGCTTTATGAGTATTTTACGGTTTATAACGAA
TTGACAAAGGTCAAATATGTTACTGAAGGAATGCGAAAACCAGCATTTCTTTCAGGTGAACAGA
AGAAAGCCATTGTTGATTTACTCTTCAAAACAAATCGAAAAGTAACCGTTAAGCAATTAAAAGA
AGATTATTTCAAAAAAATAGAATGTTTTGATAGTGTTGAAATTTCAGGAGTTGAAGATAGATTT
AATGCTTCATTAGGTACCTACCATGATTTGCTAAAAATTATTAAAGATAAAGATTTTTTGGATA
ATGAAGAAAATGAAGATATCTTAGAGGATATTGTTTTAACATTGACCTTATTTGAAGATAGGGA
GATGATTGAGGAAAGACTTAAAACATATGCTCACCTCTTTGATGATAAGGTGATGAAACAGCTT
AAACGTCGCCGTTATACTGGTTGGGGACGTTTGTCTCGAAAATTGATTAATGGTATTAGGGATA
AGCAATCTGGCAAAACAATATTAGATTTTTTGAAATCAGATGGTTTTGCCAATCGCAATTTTAT
GCAGCTGATCCATGATGATAGTTTGACATTTAAAGAAGACATTCAAAAAGCACAAGTGTCTGGA
CAAGGCGATAGTTTACATGAACATATTGCAAATTTAGCTGGTAGCCCTGCTATTAAAAAAGGTA
TTTTACAGACTGTAAAAGTTGTTGATGAATTGGTCAAAGTAATGGGGCGGCATAAGCCAGAAAA
TATCGTTATTGAAATGGCACGTGAAAATCAGACAACTCAAAAGGGCCAGAAAAATTCGCGAGAG
CGTATGAAACGAATCGAAGAAGGTATCAAAGAATTAGGAAGTCAGATTCTTAAAGAGCATCCTG
TTGAAAATACTCAATTGCAAAATGAAAAGCTCTATCTCTATTATCTCCAAAATGGAAGAGACAT
GTATGTGGACCAAGAATTAGATATTAATCGTTTAAGTGATTATGATGTCGATgcCATTGTTCCA
CAAAGTTTCCTTAAAGACGATTCAATAGACAATAAGGTCTTAACGCGTTCTGATAAAAATCGTG
GTAAATCGGATAACGTTCCAAGTGAAGAAGTAGTCAAAAAGATGAAAAACTATTGGAGACAACT
TCTAAACGCCAAGTTAATCACTCAACGTAAGTTTGATAATTTAACGAAAGCTGAACGTGGAGGT
TTGAGTGAACTTGATAAAGCTGGTTTTATCAAACGCCAATTGGTTGAAACTCGCCAAATCACTA
AGCATGTGGCACAAATTTTGGATAGTCGCATGAATACTAAATACGATGAAAATGATAAACTTAT
TCGAGAGGTTAAAGTGATTACCTTAAAATCTAAATTAGTTTCTGACTTCCGAAAAGATTTCCAA
TTCTATAAAGTACGTGAGATTAACAATTACCATCATGCCCATGATGCGTATCTAAATGCCGTCG
TTGGAACTGCTTTGATTAAGAAATATCCAAAACTTGAATCGGAGTTTGTCTATGGTGATTATAA
AGTTTATGATGTTCGTAAAATGATTGCTAAGTCTGAGCAAGAAATAGGCAAAGCAACCGCAAAA
TATTTCTTTTACTCTAATATCATGAACTTCTTCAAAACAGAAATTACACTTGCAAATGGAGAGA
TTCGCAAACGCCCTCTAATCGAAACTAATGGGGAAACTGGAGAAATTGTCTGGGATAAAGGGCG
AGATTTTGCCACAGTGCGCAAAGTATTGTCCATGCCCCAAGTCAATATTGTCAAGAAAACAGAA
GTACAGACAGGCGGATTCTCCAAGGAGTCAATTTTACCAAAAAGAAATTCGGACAAGCTTATTG
CTCGTAAAAAAGACTGGGATCCAAAAAAATATGGTGGTTTTGATAGTCCAACGGTAGCTTATTC
AGTCCTAGTGGTTGCTAAGGTGGAAAAAGGGAAATCGAAGAAGTTAAAATCCGTTAAAGAGTTA
CTAGGGATCACAATTATGGAAAGAAGTTCCTTTGAAAAAAATCCGATTGACTTTTTAGAAGCTA

AAGGATATAAGGAAGTTAAAAAAGACTTAATCATTAAACTACCTAAATATAGTCTTTTTGAGTT
AGAAAACGGTCGTAAACGGATGCTGGCTAGTGCCGGAGAATTACAAAAAGGAAATGAGCTGGCT
CTGCCAAGCAAATATGTGAATTTTTTATATTTAGCTAGTCATTATGAAAAGTTGAAGGGTAGTC
CAGAAGATAACGAACAAAAACAATTGTTTGTGGAGCAGCATAAGCATTATTTAGATGAGATTAT
TGAGCAAATCAGTGAATTTTCTAAGCGTGTTATTTTAGCAGATGCCAATTTAGATAAAGTTCTT
AGTGCATATAACAAACATAGAGACAAACCAATACGTGAACAAGCAGAAAATATTATTCATTTAT
TTACGTTGACGAATCTTGGAGCTCCCGCTGCTTTTAAATATTTTGATACAACAATTGATCGTAA
ACGATATACGTCTACAAAAGAAGTTTTAGATGCCACTCTTATCCATCAATCCATCACTGGTCTT
TATGAAACACGCATTGATTTGAGTCAGCTAGGAGGTGACTAACtcgagtaaggatctccaggca
tcaaataaaacgaaaggctcagtcgaaagactgggcctttcgttttatctgttgtttgtcggtg
aacgctctctactagagtcacactggctcaccttcgggtgggcctttctgcgtttata

**MCP-SoxS**

J23107_**Buj**_MCP-SoxS_BBa-B1002
tttacggctagctcagccctaggtattatgctagc**GAATTCATTAAAGAGGAGAAAGGTACC**at
ggggcccgcttctaactttactcagttcgttctcgtcgacaatggcggaactggcgacgtgact
gtcgccccaagcaacttcgctaacgggatcgctgaatggatcagctctaactcgcgttcacagg
cttacaaagtaacctgtagcgttcgtcagagctctgcgcagaatcgcaaatacaccatcaaagt
cgaggtgcctaaaggcgcctggcgttcgtacttaaatatggaactaaccattccaattttcgcc
acgaattccgactgcgagcttattgttaaggcaatgcaaggtctcctaaaagatggaaacccga
ttccctcagcaatcgcagcaaactccggcatctacGGTGGCGGAGGTAGCATGTCCCATCAGAA
AATTATTCAGGATCTTATCGCATGGATTGACGAGCATATTGACCAGCCGCTTAACATTGATGTA
GTCGCAAAAAAATCAGGCTATTCAAAGTGGTACTTGCAACGAATGTTCCGCACGGTGACGCATC
AGACGCTTGGCGATTACATTCGCCAACGCCGCCTGTTACTGGCCGCCGTTGAGTTGCGCACCAC
CGAGCGTCCGATTTTTGATATCGCAATGGACCTGGGTTATGTCTCGCAGCAGACCTTCTCCCGC
GTTTTCGCGCGGCAGTTTGATCGCACTCCCGCGGATTATCGCCACCGCCTGTAAGCGGCCGCca
cgcaaaaaaccccgcttcggcggggtttttcgc

**dCas9 gRNA**

>J23105_**spacer**-tracrRNA-MS2_TrrnB
tttacggctagctcagtcctaggtactatACTAGT**NNNNNNNNNNNNNNNNNNNN**GTTTTAGAG
CTAGAAATAGCAAGTTAAAATAAGGCTAGTCCGTTATCAACTTGAAAAAGTGGCACATGAGGAT
CACCCATGTGCTTTTTTTGAAGCTTGGGCCCGAACAAAAACTCATCTCAGAAGAGGATCTGAAT
AGCGCCGTCGACCATCATCATCATCATCATTGAGTTTAAACGGTCTCCAGCTTGGCTGTTTTGG
CGGATGAGAGAAGATTTTCAGCCTGATACAGATTAAATCAGAACGCAGAAGCGGTCTGATAAAA
CAGAATTTGCCTGGCGGCAGTAGCGCGGTGGTCCCACCTGACCCCATGCCGAACTCAGAAGTGA
AACGCCGTAGCGCCGATGGTAGTGTGGGGTCTCCCCATGCGAGAGTAGGGAACTGCCAGGCATC
AAATAAAACGAAAGGCTCAGTCGAAAGACTGGGCCTTTCGTTTTATCTGTTGTTTGTCGGTGAA
CT

**dCas13**

>J23107_**rc2-rbs**_dCas13_dblTerm

tttacggctagctcagccctaggtattatgctagc**GATATACTAAAAGAGGAGAAAACTGAT**AT
Gatcgaaaaaaaaaagtccttcgccaagggcatgggcgtgaagtccacactcgtgtccggctcc
aaagtgtacatgacaaccttcgccgaaggcagcgacgccaggctggaaaagatcgtggagggcg
acagcatcaggagcgtgaatgagggcgaggccttcagcgctgaaatggccgataaaaacgccgg
ctataagatcggcaacgccaaattcagccatcctaagggctacgccgtggtggctaacaaccct
ctgtatacaggacccgtccagcaggatatgctcggcctgaaggaaactctggaaaagaggtact
tcggcgagagcgctgatggcaatgacaatatttgtatccaggtgatccataacatcctggacat
tgaaaaaatcctcgccgaatacattaccaacgccgcctacgccgtcaacaatatctccggcctg
gataaggacattattggattcggcaagttctccacagtgtatacctacgacgaattcaaagacc
ccgagcaccatagggccgctttcaacaataacgataagctcatcaacgccatcaaggcccagta
tgacgagttcgacaacttcctcgataaccccagactcggctatttcggccaggcctttttcagc
aaggagggcagaaattacatcatcaattacggcaacgaatgctatgacattctggccctcctga
gcggactggcgcactgggtggtcgctaacaacgaagaagagtccaggatctccaggacctggct
ctacaacctcgataagaacctcgacaacgaatacatctccaccctcaactacctctacgacagg
atcaccaatgagctgaccaactccttctccaagaactccgccgccaacgtgaactatattgccg
aaactctgggaatcaaccctgccgaattcgccgaacaatatttcagattcagcattatgaaaga
gcagaaaaacctcggattcaatatcaccaagctcagggaagtgatgctggacaggaaggatatg
tccgagatcaggaaaaatcataaggtgttcgactccatcaggaccaaggtctacaccatgatgg
actttgtgatttataggtattacatcgaagaggatgccaaggtggctgccgccaataagtccct
ccccgataatgagaagtccctgagcgagaaggatatctttgtgattaacctgaggggctccttc
aacgacgaccagaaggatgccctctactacgatgaagctaatagaatttggagaaagctcgaaa
atatcatgcacaacatcaaggaatttaggggaaacaagacaagagagtataagaagaaggacgc
ccctagactgcccagaatcctgcccgctggccgtgatgtttccgccttcagcaaactcatgtat
gccctgaccatgttcctggatggcaaggagatcaacgacctcctgaccaccctgattaataaat
tcgataacatccagagcttcctgaaggtgatgcctctcatcggagtcaacgctaagttcgtgga
ggaatacgcctttttcaaagactccgccaagatcgccgatgagctgaggctgatcaagtccttc
gctagaatgggagaacctattgccgatgccaggagggccatgtatatcgacgccatccgtattt
taggaaccaacctgtcctatgatgagctcaaggccctcgccgacacctttttccctggacgagaa
cggaaacaagctcaagaaaggcaagcacggcatgagaaatttcattattaataacgtgatcagc
aataaaaggttccactacctgatcagatacggtgatcctgcccacctccatgagatcgccaaaa
acgaggccgtggtgaagttcgtgctcggcaggatcgctgacatccagaaaaaacagggccagaa
cggcaagaaccagatcgacaggtactacgaaacttgtatcggaaaggataagggcaagagcgtg
agcgaaaaggtggacgctctcacaaagatcatcaccggaatgaactacgaccaattcgacaaga
aaaggagcgtcattgaggacaccggcagggaaaacgccgagagggagaagtttaaaaagatcat
cagcctgtacctcaccgtgatctaccacatcctcaagaatattgtcaatatcaacgccaggtac
gtcatcggattccattgcgtcgagcgtgatgctcaactgtacaaggagaaaggctacgacatca
atctcaagaaactggaagagaagggattcagctccgtcaccaagctctgcgctggcattgatga
aactgcccccgataagagaaaggacgtggaaaaggagatggctgaaagagccaaggagagcatt
gacagcctcgagagcgccaaccccaagctgtatgccaattacatcaaatacagcgacgagaaga
aagccgaggagttcaccaggcagattaacagggagaaggccaaaaccgccctgaacgcctacct

gaggaacaccaagtggaatgtgatcatcagggaggacctcctgagaattgacaacaagacatgt
accctgttcgcaaacaaggccgtcgccctggaagtggccaggtatgtccacgcctatatcaacg
acattgccgaggtcaattcctacttccaactgtaccattacatcatgcagagaattatcatgaa
tgagaggtacgagaaaagcagcggaaaggtgtccgagtacttcgacgctgtgaatgacgagaag
aagtacaacgataggctcctgaaactgctgtgtgtgcctttcggctactgtatccccaggttta
agaacctgagcatcgaggccctgttcgataggaacgaggccgccaagttcgacaaggagaaaaa
gaaggtgtccggcaattcctaactcgagtaaggatctccaggcatcaaataaaacgaaaggctc
agtcgaaagactgggcctttcgttttatctgttgtttgtcggtgaacgctctctactagagtca
cactggctcaccttcgggtgggcctttctgcgtttata

**dCas13 gRNA**

>J23110_Direct-repeat_**spacer**_ECK120033736
tttacggctagctcagtcctaggtacaatgctagcCACTAGTGCGAATTTGCACTAGTCTAAAA
C**NNNNNNNNNNNNNNNNNNNNNNNNNN**AAAGCCCCGGAAGATCACCTTCCGGGGCTTTtt
tattgcgc

**LNT pathway**

>J3_J23117_**RBS-A**_lacY
AGCATTTGCGATCATTCACGCAGCGCTTATTCAGTTGCTCACTGCGATGTCATAATCATCGCTA
CGAGCTGTGAAAGATGCATAAAGCTCGTACGACGCGTTCGCTCGTCTCCTCACTTCTCCTACGG
AGCGTTCTGGACACAACGTCGTCTTGAAGTTGCGATTATAGAttgacagctagctcagtcctag
ggattgtgctagc**AAAGATCTTTTAAGAAGGAGATATACAT**atgtactatttaaaaaacacaaa
cttttggatgttcggtttattctttttcttttactttttatcatgggagcctacttcccgttt
ttcccgatttggctacatgacatcaaccatatcagcaaaagtgatacgggtattatttttgccg
ctatttctctgttctcgctattattccaaccgctgtttggtctgctttctgacaaactcgggct
gcgcaaatacctgctgtggattattaccggcatgttagtgatgtttgcgccgttctttattttt
atcttcgggccactgttacaatacaacattttagtaggatcgattgttggtggtatttatctag
gcttttgttttaacgccggtgcgccagcagtagaggcatttattgagaaagtcagccgtcgcag
taatttcgaatttggtcgcgcgcggatgtttggctgtgttggctgggcgctgtgtgcctcgatt
gtcggcatcatgttcaccatcaataatcagtttgttttctggctgggctctggctgtgcactca
tcctcgccgttttactctttttcgccaaaacggatgcgccctcttctgccacggttgccaatgc
ggtaggtgccaaccattcggcatttagccttaagctggcactggaactgttcagacagccaaaa
ctgtggtttttgtcactgtatgttattggcgtttcctgcacctacgatgtttttgaccaacagt
ttgctaatttctttacttcgttctttgctaccggtgaacagggtacgcgggtatttggctacgt
aacgacaatgggcgaattacttaacgcctcgattatgttctttgcgccactgatcattaatcgc
atcggtgggaaaaacgccctgctgctggctggcactattatgtctgtacgtattattggctcat
cgttcgccacctcagcgctggaagtggttattctgaaaacgctgcatatgtttgaagtaccgtt
cctgctggtgggctgctttaaatatattaccagccagtttgaagtgcgttttcagcgacgatt
tatctggtctgtttctgcttctttaagcaactggcgatgatttttatgtctgtactggcgggca
atatgtatgaaagcatcggtttccagggcgcttatctggtgctgggtctggtggcgctgggctt

caccttaatttccgtgttcacgcttagcggccccggcccgctttccctgctgcgtcgtcaggtg
aatgaagtcgcttaa

>J5_J23117_**RBS-B**_lgtA
TGGCAATTCCGACGTCTATACATCGCATCACTACACTATTGATTATCATTGTGTACGTAACGAG
CTTGCACAACGTGAAGTTCTTCGAGCACTTCAGCTCGCAACGTAAATGACAGTTGCTGTTAAGT
GACGTGAATCCTTCAATGCTGCTCATGCTGCTGTCGTAAATAAGTAAGTCACTCCCACttgaca
gctagctcagtcctagggattgtgctagc**AACTGGTAATTTGAGGAGGTAATTT**ATGCCGTCGG
AAGCCTTTCGTCGCCATCGTGCGTACCGTGAAAACAAACTTCAGCCTTTGGTGTCTGTTTTGAT
CTGCGCGTATAATGTAGAGAAGTATTTTGCACAGAGTCTGGCTGCGGTTGTGAACCAAACCTGG
CGTAACTTAGACATCTTAATCGTCGATGACGGAAGTACAGATGGCACGTTGGCGATCGCACAAC
GTTTCCAAGAGCAAGATGGTCGCATCCGTATCTTAGCCCAGCCACGCAATTCGGGACTTATCCC
ATCTTTGAACATTGGACTTGATGAACTGGCTAAAAGTGGCATGGGAGAATACATTGCGCGCACT
GATGCGGATGACATTGCAGCGCCCGACTGGATCGAGAAAATCGTGGGAGAGATGGAGAAAGATC
GCTCTATCATCGCTATGGGAGCATGGTTAGAGGTTCTGTCAGAGGAGAAGGACGGCAACCGTTT
AGCTCGTCACCATGAGCATGGAAAAATCTGGAAAAAACCCACCCGCCACGAAGACATCGCCACG
GTCTTCCCATTCGGTAACCCAATCCACAATAATACAATGATTATGCGTCGTAGCGTGATTGATG
GAGGTCTTCGCTACAACACTGAGCGCGATTGGGCCGAGGATTACCAGTTTTGGTATGACGTGAG
CAAGTTAGGCCGTCTGGCTTACTATCCAGAAGCGCTGGTGAAGTACCGTCTTCATGCGAACCAG
GTGTCAAGTAAATACAGTGTACGTCAACATGAAATCGCACAGGGTATCCAAAAAACAGCGCGCA
ATGATTTTTTGCAGAGCATGGGTTTTAAAACACGCTTCGACAGCTTGGAGTATCGCCAAATCAA
AGCCGTAGCCTATGAATTACTGGAAAAGCATCTGCCGGAGGAAGATTTTGAGCGTGCCCGCCGC
TTTTTATACCAATGTTTTAAACGCACCGACACCCCACCAGCTGGCGCCTGGCTTGATTTTGCCG
CCGACGGCCGTATGCGTCGCTTATTTACTTTGCGCCAATATTTCGGTATCTTACGCCGTCTGTT
GAAAAATCGCTGA

>J6_J23117_**RBS-C**_galT
cctgcacgagttcgctgtcgagacaagtctcttagcgacgtattacgaagatcacatagtcaga
tgaagctatagagcacgacgctaacgattacgtcacgcttgacacaacagtttcgctacctagt
gctcgcgcgactgcgacgttgtccttctagtcgcccatgactcttgacagctagctcagtccta
gggattgtgctagc**AATCTCATAAATCAAATATAGGGAGGATCAT**ATGGACACCATCATGATTA
AACGTCCGCTGGTTAGCGTTATTCTGCCGGTGAATAAAAACAATCCGCATCTGGAAGAAGCAAT
CCAGAGCATTAAAAACCAGACCTATAAAGAGCTGGAACTGATCATTATTGCCAACAACTGCGAG
GATAACTTTTATAGCCTGCTGCTGAAATATCAGGACCAGAAAACCAAAATTATCCGCACCAGCA
TCAAATATCTGCCGTTTAGCCTGAATCTGGGTGTTCATCTGAGCCAGGGTGAATATATTGCACG
TATGGATTCAGATGATATCAGCGTTCTGGATCGCATTGAAAAACAGGTTAAACGCTTTCTGAAT
ACACCGGAACTGAGCATTCTGGGTAGCAATGTTGAATATATCAATGAAGCCAGCGAAAGCATTG
GCTATAGCAACTATCCGCTGGATCATAGCAGCATTGTTAATAGCTTTCCGTTTCGTTGTAATCT
GGCACATCCGACCATTATGGTTAAAAAAGAAGTGATTACCACGCTTGGTGGCTATATGTATGGT
AGCCTGAGCGAAGATTATGATCTGTGGATTCGTGCAAGCCGTCATGGCAATTTCAAATTTAGCA
ATATTGATGAACCGCTGCTGAAGTACCGTATTCATAAAGGTCAGGCAACCAATAAAAGCAACGC

CTATAACATCTTTGCCTTTGATAGCAGCCTGAAAATCCGTGAATTTCTGCTGAATGGTAATGTG
CAGTATCTGCTGGGTGCAGCACGTGGTTTTTTTGCATTTCTGTATGTGCGCTTCATCAAAAAAT
GA
